# *Drp1-JNK* knockdown mitigates Scribble loss induced cell proliferation, metastasis and lethality phenotypes in *Drosophila*

**DOI:** 10.1101/2024.04.10.588977

**Authors:** Jyotsna Singh, Saripella Srikrishna

**Author notes:** Address for correspondence: Saripella Srikrishna, Ph.D. Cancer and Neurobiology Laboratory Department of Biochemistry, Institute of Science Banaras Hindu University, Varanasi 221005, Uttar Pradesh, India.

## Abstract

Mitochondrial dynamics are emerging as master regulators for targeting several types of cancers, including breast cancer, cervical cancer, and hepatocellular carcinoma, for therapeutic intervention. Mitochondrial morphology, size, position and activity within cells is regulated by dynamic fission and fusion events. Dynamin-related protein 1 (*Drp1*) promotes mitochondrial fission and maintains mitochondrial homeostasis. Loss of *Scrib* is implicated in several human cancers wherein mitochondrial dysfunction leads to excessive cell proliferation and metastasis. However, the exact molecular mechanisms behind the *Scrib* loss induced dysregulation of mitochondrial dynamics in cancer progression remains obscure. Although the role of mitochondrial dynamics are being investigated in several types of cancers, but the role of *Drp1*- mediated fission event in regulating the maintenance of polarity of cells upon loss of *Scrib* function is elusive. In this study, for the first time, we blocked the function of *Drp1* activity in *Scrib* knockdown induced metastasis cancer model by two ways, firstly, through genetic ablation of *Drp1,* and secondly by using mdivi-1, a *Drp1* specific inhibitor. Genetic depletion of *Drp1* expression (*Drp1^RNAi^*) in *Scrib* knockdown cells inhibits Metalloproteinase *MMP1*, reduces ROS production, restores apico-basal (A/B) cell polarity and enhances ATP production. Further to confirm role of Drp1 in regulation of cell polarity, we employed mdivi, a Drp1 specific inhibitor which has dose dependent effect in cell polarity regulation. This study also reveals that *JNK* inhibition (*JNK^RNAi^*) in *Scrib* abrogated cells mitigates the *Drp1* expression and controls cell proliferation leading to restoration of mitochondrial morphology and epithelial cellpolarity. Our results highlight *Drp1* as a key regulator in maintaining the apico-basal polarity of cells which gets affected upon loss of *Scrib* but *Drp1-JNK* downregulation effectively mitigates *Scrib^RNAi^* associated cell proliferation, metastasis and pupal lethality phenotypes.

## Introduction

Scribble (*Scrib*), is a tumor suppressor gene and apico-basal (A/B) polarity regulator in epithelial cells [1]. Many cancers of epithelial origin like cervical cancer [2], breast cancer [3,4], ovarian cancer, colorectal cancer, pituitary tumor etc., havebeen reported to exhibit mitochondrial dysfunction [5] but the underlying mechanismremains elusive. Mitochondrial disruption is the major source of increasedreactive oxygen species(ROS)production [6]. Loss of *Scrib* is associated with ROS enhancement in cancer cells, however, the regulatory mechanisms are largely unknown [7]. According to several studies, abnormal cell proliferation needs excessive production of ROS in order to improve cancer cell growth by an oncogenic cooperative signal [3,8].

In normal cells, the dynamic behavior of mitochondria isimportant for maintenance of cellular homeostasis [9]. The dynamics of mitochondrialmembrane machinery maintain the balance between the mitochondrial fission and fusion transition events [10]. The fusion of outer mitochondrial membrane (OMM) is regulated by mitofusin/Marf (Mitochondrial Assembly Regulatory Factor) in *Drosophila*, and Mfn (mitofusin) in mammals while inner mitochondrial membrane (IMM) fusion is regulated by optic atrophy 1 (OPA1) and mitochondrial fission is regulated by dynamin-related protein 1 (*Drp1*). These two mitochondrial regulators encode a dynamin-family GTPase, to maintain the mitochondrial morphology and activity [11]. The *Drp1* migrates from cytosol to mitochondrial outer membrane (MOM) and forms anoligomeric collar like structure around the mitochondrion and hydrolyzes GTP to execute *Drp1*-mediated mitochondrial fission event [12, 13].

In the current study, we have explored the role of *Drp1* in tumor progression upon knockdown of *Scrib* in *Drosophila*. Our previous studies have shown that upregulation of *Drp1* expression leads to mitochondrial dysfunction in *Scrib* knockdown tumor cells [14]. However, so far, there are no reports available, that demonstrate that *in vivo* blocking of *Drp1* expressionin the background of *Scrib*loss of function are associated with cancer cell proliferation in *Drosophila*. For the first time we show that knockdown of *Drp1* expression in *Scrib* knockdown cells results in restoration of cell polarity and rescues the pupal death phenotype. Moreover, to further determine the role of *Drp1* in cancer cell migration and invasion upon *Scrib* knockdown, we employed Mdivi (Mitochondrial division inhibitor), a known inhibitor of *Drp1,* which inhibits its GTPase activity [15]. Our results shows Mdivi-1 induces a dose-dependent inhibition on cancer cell migration and restores the cell polarity upon loss of *Scrib*.

In *Drosophila*, well conserved *JNK* (c-Jun N-terminal kinase) signaling pathway activation promotes cell proliferation upon *Ras* activation and cooperative association with cell polarity regulator genes [16, 17]. *JNK* pathway’s activation leads to mitochondrial dysfunction and excess ROS production due to the translocation of JNK protein on to the outer membrane of mitochondria in cancer cells [18,19]. Previous studies report that *JNK* mediates *Drp1* translocation to regulate mitochondrial fission event [20]. In this context, our studies show that *Scrib* regulates *Drp1/JNK* activity to maintain polarity of cells and regulates mitochondrial dynamics. To the best of our understanding, this study reveals a new insight into the mechanism for maintaining mitochondrial morphology and A/B cell polarity via the *Drp1* dependent mitochondrial fission in *Scrib* abrogated tumor cells.

## Materials and Methods

### Fly stocks and culture conditions

We used Oregon-R +/+ as a Wild type, UAS-*Scrib^RNAi^*(#35748),UAS-GFP (#1521), UAS- *Drp1^RNAi^* (#51483), UAS-*Hep^RNAi^* (#2190-R2), Sp/Cyo; dCO2/TM6B, mitocherry Red OMM; Ptc-Gal4 and Ptc-GAL4 (#2017) fly strains for this study. The Ptc-gal4 driver linedrives the expression of the transgene at the anterior/posterior (A/P) boundary region of *Drosophila* wing imaginal discs. Flies strain were reared on corn agar medium that contained in 1000ml water: 5.6g of agar, 47.2g of Maize powder, 41.67g of sugar, 16.7g of yeast, 6.94ml absolute ethanol solution used to dissolve 2.8g of nepagin (anti-fungal) and 2.8ml of propionic acid (anti- bacterial). Fly stocks were obtained from the Bloomington Drosophila Stock Center (https://flybase.bio.india na.edu). All crosses were carried out at 24^0^C in standard corn-meal agar media and fly stocks were maintained in B.O.D incubator.

### Wing disc area quantification

The roaming third instar larvae of Wild type, Scribble knockdown (*Scribble^RNAi^*) and rescued genotype (*UAS-Drp1^RNAi^; UAS-Scrib^RNAi^* and *UAS-Hep^RNAi^; UAS-Scrib^RNAi^*), were dissected in 1X PBS on cavity slide and separated wing imaginal discs which were transferred to the maximo cavity slide. Then the discs were fixed in 4% paraformaldehyde for 15 minutes at room temperature, followed by washing it with 1XPBST (0.1%), 3 times for 5 min each. The discs were mounted on glass slides with DABCO respectively. Bright flied images were obtained by confocal microscope (Zeiss LSM-510 Meta). To measure wing disc area using open free shape curve drawing mode in overlay option in offline LSM software (indicate by yellow line). The histogram represents average wing imaginal area disc for each genotype.

### Generation of transgenic flies

Different crosses were set to bring *Drp1^RNAi^* and *Hep^RNAi^* in genetic combination (introgression) together with Scrib*^RNAi^*. Also, to monitor tumor progression in wing imaginal disc, we genetically labeled *Scrib* knockdown cells using wing discs specific Ptc-Gal4 with a visible marker such as green fluorescent protein (GFP) (Supplementary Information, Scheme 1-4).

### Wing Venation pattern analysis

Rescued adult flies were anaesthetized and the wings from the thorax region were carefully detached with the help of sharp dissection needles. The isolated wing was placed on the top of slide, protected with a coverslip, the edges of which were completely sealed with a transparent nail polish. The images for wing venation pattern were captured under bright-field filter using fluorescence Nikon-NiU microscope. The length of L2 to ACV, length of ACV that bridge L3- L4 and PCV that bridge L4-L5 is measured with offline NIS-element BR 4.3 software for alteration analysis.

### Lifespan Assay

The lifespan of adult rescue flies of *UAS-Drp1^RNAi^; UAS-Scrib^RNAi^* and *UAS-Hep^RNAi^; UAS- Scrib^RNAi^*genotypes were measured from the day of eclosion at 24°C. A minimum of 100 flies were taken (10 flies/vial) for each genotype. The total number of surviving flies were checked and counted every day. The flies were transferred in fresh food vials every 2 days, the number of dead flies was scored for each genotypes and survival curve generated by using Graph pad prism 5 software.

### Drug screening in Cancer bearing larvae

To test whether *Drp1* inhibition positively regulates the polarity of cells, we tested the effect of small chemical molecule *Drp1* inhibitor, mdivi on *Scrib* knockdown tumor bearing larvae. The mdivi was obtained from Sigma-Aldrich (M0199), pre-dissolved in DMSO (dimethylsulfoxide) and mixed in corn agar food medium. 100% DMSO shows toxic effect on flies, so, we prepared drug solution in 0.3 % DMSO considered as no observed adverse effect level (NOAEL), non-toxic concentration used for the *in-vivo* drug screening in *Drosophila melanogaster*[21,22]. Control flies treated with DMSO and without DMSO were used for the experiment. The cancer bearing larvae were fed with drug treated food from higher to lower concentration. Drug treatment was analyzed at six different concentrations, 15µM, 30µM, 60µM, 120µM, 250µM and 500µM to test dose dependent effect tumor development. As 15µM and 30µM drug treated larvae, did not show any phenotypic variation in tumor growth, so further experimental analysis where performed at above 30µM drug doses.

### Immunostaining of larval wing imaginal discs

The *Drosophila* third instar larval wing imaginal discs were isolated from the desired genotypes. Dissection was performed in 1X PBS (phosphate buffer saline) solution. Tissue were fixed in 4% paraformaldehyde (PFA) for 15 min at RT. The wing discs were washed in 0.2% PBST (1X PBS, 0.2% Triton-X) three times, 5 min each and tissue were blocked in blocking solution (3% bovine serum albumin (BSA) solution in 1X PBS) for 1hr at RT followed by incubation with the required primary antibodyin blocking solution for overnight at 4°C. A cocktail of three mouse anti-MMP1 (matrix metalloproteinases-1) catD monoclonal antibodies raised against the catalytic domain (14A3D2, 3B8D12, 5H7B11 used in 1:1:1 dilution) was obtained from the developmental study hybridoma bank (DSHB), Iowa. Rabbit anti-Drp1 (FITC-DRP1 used 1:100 dilution) was obtained from the FABGENIX, Rabbit anti-ACTIVE JNK pAb, (v7931 used in 1:1000 dilution) from Promega and Rabbit anti-Ph3 was obtained from Merck Millipore (06-570 used in 1:1000 dilution). The tissues were washed with 0.2% PBST three times, 5 min each, followed by incubation with the desired secondary antibodyin 1X PBS for 3hrs at RT. Secondary antibodies used were, goat anti-mouse IgG, Alexa Fluor 594 (1:1000 dilution, Cat# A-11005) and goat anti-rabbit IgG, Alexa Fluor 488 (1:1000 dilution, Cat# A-11008). Tissues were washed with 0.1% PBST three times, 5 min each following the 2^0^ incubation, further incubated with DAPI (1µg/ml in 1X PBS) for 10min then again washed with 0.1% 1X PBST (1X PBS, 0.1% Triton-X) three times, 5 min each and then sample were mounted in DABCO. Immunofluorescence images were taken using confocal microscope (Zeiss LSM-510 Meta) and processed using LSM software and arranged in Adobe Photoshop 7.0.

### RNA isolation and cDNA synthesis

Total RNA was extracted from desired genotype (Wild type (Oregon R^+)^, *Scribble^RNAi^* and rescued genotype (*UAS-Drp1^RNAi^; UAS-Scrib^RNAi^* and *UAS-Hep^RNAi^; UAS-Scrib^RNAi^*), using TRI reagent (TAKARA) as per manufacturer’s protocol. The quantity of RNA was checked on a 1% agarose gel and documented using Gel Doc (BIO-RAD). High quantity RNA from both samples estimated by absorbance A260/280 ratio was reverse transcribed to cDNA. For cDNA preparation 1µg of RNA, 1µl random hexamer (Applied Biosystems) and 11µl (0.1%) DEPC treated water was run for single cycle at 70^0^C for 5min. in a thermal cycler (BIO-RAD). To this 2µl 10mM dNTPs (Invitrogen) and 0.5µl (200U/µl) RNase inhibitor (Applied Biosystems) were added and run for single cycle at 25^0^C for 5 min. Finally, 1µl (100U/µl) reverse transcriptase (Ambion) was added and whole mixture was run at 37^0^C for 1h in a thermal cycler (Sure Cycler 8800, Agilent).

### Qualitative Real-time PCR (qRT-PCR)

Prepared cDNA of desired genotype was used as a template for quantification of differentially expressed genes, normalized against GAPDH with specific primers (Table.1) using SYBR (R) GREEN JUMPSTART TAQ Ready mix (Thermo Fisher Scientific) using Real-time thermal cycler analysis (ΔCT values).

**Table. 1.**
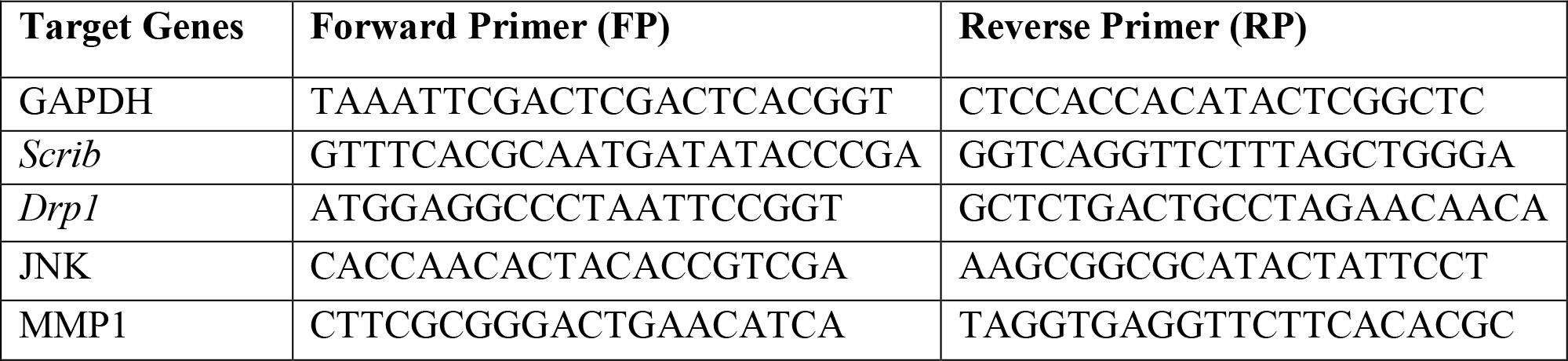
Specific primers (shown in 5’-3’ direction) for different genes.

### Western blotting

Total protein was extracted using lysis buffer and protein quantification was done with Bradford assay. Equal amount of protein from defined experimental groups were loaded and resolved on the 10% SDS-PAGE,wet transferred onto PVDF membrane at 4°C. Blot was probed with primary antibodies with overnight incubation. The antibodies used are, a cocktail of three monoclonal mouse antibodies for MMP1 (1:100, DSHB, Cat No. 3A6B4, 3B8D12 and 5H7B11), polyclonal rabbit antibody for Drp1 (1:500, Cat No. DRP1 FITC), and monoclonal mouse antibody for β-tubulin (1:125, Cat No.DSHB-S1-810-(DSHB)-E7 anti beta tubulin supernatant 1ml).Proteins were detected using Goat Anti-Mouse IgG (H+L) peroxidase Conjugates (1;2500, Thermo scientific, Cat No. 31430) and Goat Anti-Rabbit IgG-HRP (1;1000, Merck, Cat No. 032102). The specific bands were detected using the ECL (Enhanced chemiluminescence detection, Cat No. 1705060 (Genetix)-Clarity Western ECL Subs, 200ml) system. Gene expression was analyzed after normalizing β-tubulin expression using image J softwere.

### In silico Analysis

Protein-Protein interaction signaling pathways were analyzed by STRING 9.0 web software. STRING is a functional protein connection networks analysis (https://string-db.org) using protein accession number to analyzed the connecting link between molecular signaling pathways and interaction associated between the differential expressed proteins [23].

### ROS estimation

To detect in-vivo intracellular ROS production in desired wing imaginal disc, we used H2DCFDA (2’,7’-dichlorodihydro fluorescein diacetate, Sigma, D6883) staining. H2DCFDA is a cell membrane permeable indicator used for detection of ROS. Oxidation of H2DCFDA, when it reacts with H2O2, generates highly green fluorescent 2’,7’-dichlorofluorescein (DCF). The acetate group H2DCFDA hydrolyzed by intracellular esterase converts non-fluorescent H2DCFDA into highly green fluorescent DCF [24].

Third instar larval wing imaginal discs of desired genotype were isolated in 1X PBS (pH7.2) and discs were incubated in H2DCFDA (working concentration 5µg/ml) for 30minute at 37°C, washed with 1X PBS and images were captured in fluorescence microscope and processed using software NIS-Elements (BR).

### Biochemical Assays

#### SOD Assay by NBT method

SOD assay was used for measuring the superoxide dismutase (SOD) enzyme inhibition activity. The samples of desired genotype were homogenized in 50µl extraction buffer (100mM K2HPO4, 100mM KH2PO4 and 1mM EDTA) and centrifuged at 12,000 rpm for 10 minute at 4°C. The supernatants were transferred to fresh 1.5ml micro centrifuge tubes and concentration of protein was estimated by Bradford method [25]. Further, in a clean glass tube 900µl distilled water, 200µl L-methionine (20mM) and 100µl NBT (2.25mM) (nitro blue tetrazolium),(which produce light blue color formazan after reducing the superoxide anion), and 1.5ml of extraction buffer were added for 100µg protein sample. In dark condition, sample was incubated for 30 minute under yellow light after adding 100µl riboflavin (60µM). Absorbance were taken at 560nm using multimode plate reader Synergy H1. Then to measure 50% inhibition of NBT reduction by superoxide dismutase (SOD) by subtraction the sample OD with blank OD then divide with blank OD (Blank OD-Sample OD/ Blank OD*100). 50% SOD inhibition is equal to 1 SOD unit then 1% equal to 1/50 multiple by 50% inhibition of NBT reduction. The SOD activity was represented by units per min per mg protein sample.

#### H2O2 (Hydrogen peroxide) Assay

The sample of desired genotype were homogenized in sodium phosphate buffer (50mM, pH 6.6) containing Na2HPO4 and NaH2PO4, and centrifuged at 10,000rpm for 10 minute at 4°C. The supernatants were transferred to fresh 1.5ml micro centrifuge tube and concentration of protein is estimated by Bradford method. In 100µg sample protein added 1ml TiSO4 (0.1% in 20% H2SO4) and incubated for 10 minute at room temperature. Further, it was centrifuged at 1000rpm for 20 minute at 4°C. Absorbance were taken at 410nm using multimode plate reader Synergy H1.

#### Catalase (CAT) Assay

The sample of desired genotype were homogenized in catalase extraction buffer (pH 8.0) containing Tris-HCl (50mM), EDTA (0.5mM) and 2% (w/v) PVP (polyvinyl pyrrolidone). The homogenized sample were centrifuge 10,000rpm for 10 minute at 4°C. The supernatants were transferred to fresh 1.5ml micro centrifuge tube and concentration of protein was estimated by Bradford method. The assay mixture contains 100µl crude total protein, 1ml potassium buffer (contain 100mM K2HPO4, 100mM KH2PO4) and 20mM of 8.8M H2O2 solution. The H2O2 consumption was measured 10times at 5 second interval and absorbance was recorded at 240nm. The catalase activity was calculated by subtracting highest value of absorbance to lowest value of absorbance 240nm divided by time log (5 sec) multiply by extinction coefficient 0.036.

#### TBARS (Lipid peroxidation) Assay

The Thiobarbituric acid reactive substance (TBARS) assay also known as lipid peroxidation assay is used for detection of lipid oxidation by measuring end product, the malondialdehyde (MDA). The sample of desired genotype were homogenized in homogenizing buffer (1XPBS (1ml), 10µl PMSF) and centrifuged at 10,000 rpm for 20 minute at 4°C. The supernatants were transferred to a fresh 1.5ml micro centrifuge tube and concentration of protein was estimated by Bradford method. In total extracted protein sample,1.6ml d.H2O, and 100µl SDS (10%) were added and after 5-minute incubation at RT, 20% acetic acid (600µl) was added and further incubated at RT for 2 minute. 0.8% TBA (Thiobarbituric acid) was added to the sample, mixed properly and placed in 90°C in water bath for 1hour at. Subsequently, the samples were cooled and centrifuged at 10,000rpm for 10minute at 4°C. Absorbance were taken at 532nm and 600nm using multimode plate reader Synergy H1. The MDA concentration calculation can be done by subtracting absorbance reading at 532nm to absorbance reading at 600nm divided by 155, which is extinction coefficient of MDA.

### Analysis of Mitochondrial structural morphology

Mitochondrial structure was analyzed by employing Ptc-Gal4 driver line and UAS-mitocherry red (Standard genetic cross schemes shown in Scheme. 2). Wing imaginal discs from roaming third instar larvae of UAS-mitocherry; Ptc-Gal4, *Scribble^RNAi^* (UAS-mitocherry; Ptc- Gal4<*Scribble^RNAi^*) and rescued genotype (UAS-mitocherry; Ptc-Gal4<*UAS-Drp1^RNAi^; UAS- Scrib^RNAi^*and UAS-mitocherry; Ptc-Gal4<*UAS-Hep^RNAi^; UAS-Scrib^RNAi^*), were dissected in IX PBS and were fixed in 4% paraformaldehyde for 15 minutes at room temperature, followed by washing with 1XPBST (0.1%), 3 times for 5 min each. The discs were mounted on glass slides with DABCO. Images were obtained by confocal microscope (Zeiss LSM-510 Meta).

### *In-situ* cell death detection (TUNEL) Assay

Detection of cell death was carried out using *In-situ* cell death detection kit, TMR red (Roche Diagnostics, REF 12156792910). Wing imaginal discs from roaming third instar larvae of wild type, cancer *Scribble^RNAi^*and rescued genotype (UAS-*Drp1^RNAi^; UAS-Scrib^RNAi^* and UAS- *Hep^RNAi^; UAS-Scrib^RNAi^*), were dissected in 1X PBS and were fixed in 4% paraformaldehyde for 15 minutes, followed by washing it with 1XPBST, 3 times for 5 min each for permeabilization and washed three times with 1X PBS, 5 min each followed by incubation with TUNEL (TdT-mediated dUTP-X nick end labelling) reaction mixture at 37°C for 2 hrs followed by washing with 1XPBS for two times 5 min each. The working solution of TUNEL reaction mixture were prepared immediately just before use by mixing the 50µl of enzyme solution (TdT) in 450µl of label solution (TMR-dUTP). The discs were mounted on glass slides with DABCO. Images were obtained by confocal microscope (Zeiss LSM-510 Meta).

### ATP Consumption Assay

The sample of desired genotype were homogenized in 50µl ATP assay buffer and centrifuged at 12,000 rpm for 15mintute at 4°C. The total ATP production was measure in nm/µg of protein by the ATP colorimetric assay kit (Sigma, MAK190-1KT) and experiment was performed as per provided manufacturer’s procedure. The developed pink color is stable for 2 hours. The absorbance was taken at 570nm in multimode microplate reader. The protein concentration of the sample was measure by the Bradford method and ATP level was normalized with protein content.

### Statistical Analysis

The statistical analysis was performed using Graph Pad Prism 5.0 Software Inc. All data results with error bar represented as mean ± standard error mean (SEM) used for statistical analysis for three independent experiments. The data were analyzed by using one-way ANOVA (analysis of variances) followed by Bonferroni multiple comparison test and two-tailed student’s t test for real-time PCR shows statistical difference between wildtype and tested group of indicated genotypes. Survival assay or Life span assay was performed by Kaplan-Meier method and significant was calculated by Log-rank (Mantel-Cox) test. P<0.05 was considered a statistically significance difference for data analysis.

## Results

### Knockdown of *Drp1 and JNK* signaling in *Scrib^RNAi^* cells inhibit metastasis, polarity loss and absolute pupal lethality

In previous reports, loss of *Scrib* is shown to exacerbate the cell polarity defects and disrupt cell-cell junction integrity [26]. We used the wing specific patched-Gal4 (Ptc-Gal4) to knockdown the expression of *scrib* through *UAS-Scrib^RNAi^*, specifically in anterior posterior boundary region in wing disc. We found that *scrib* knockdown leads to development of giant tumor bearing larvae (Ptc-Gal4>UAS-*Scrib^RNAi^*) (Fig.1 A and B). For a First, we show the rescue of *scrib* knockdown induced larval phenotypes upon *Drp1* depletion (Ptc-Gal4>UAS-*Drp1^RNAi^*; UAS-*Scrib^RNAi^*) (Fig. 1. C). Further,*JNK* signaling upstream regulator *Hemipterous* (*Hep*), also known as Jun Kinase Kinsae (JNKK), knockdown in *Scrib* abrogated cells (Ptc-Gal4>*UAS-Hep^RNAi^; UAS-Scrib^RNAi^*) suppresses tumor development (Fig. 1. D). The wing disc area from the *Scrib* knockdown larvae shows that the tumorous wing disc is smaller in size area wise with an increased volume compared to wild type (Fig.1I) and this phenotype is rescued upon knockdown of *Drp1* and *JNK* signaling (UAS-*Drp1^RNAi^*, UAS-*Scrib^RNAi^*and *UAS-Hep^RNAi;^ UAS-Scrib^RNAi^*) as shown in Fig.1. (E-H). The wing disc area (Fig.1.I).

**Figure 1.**
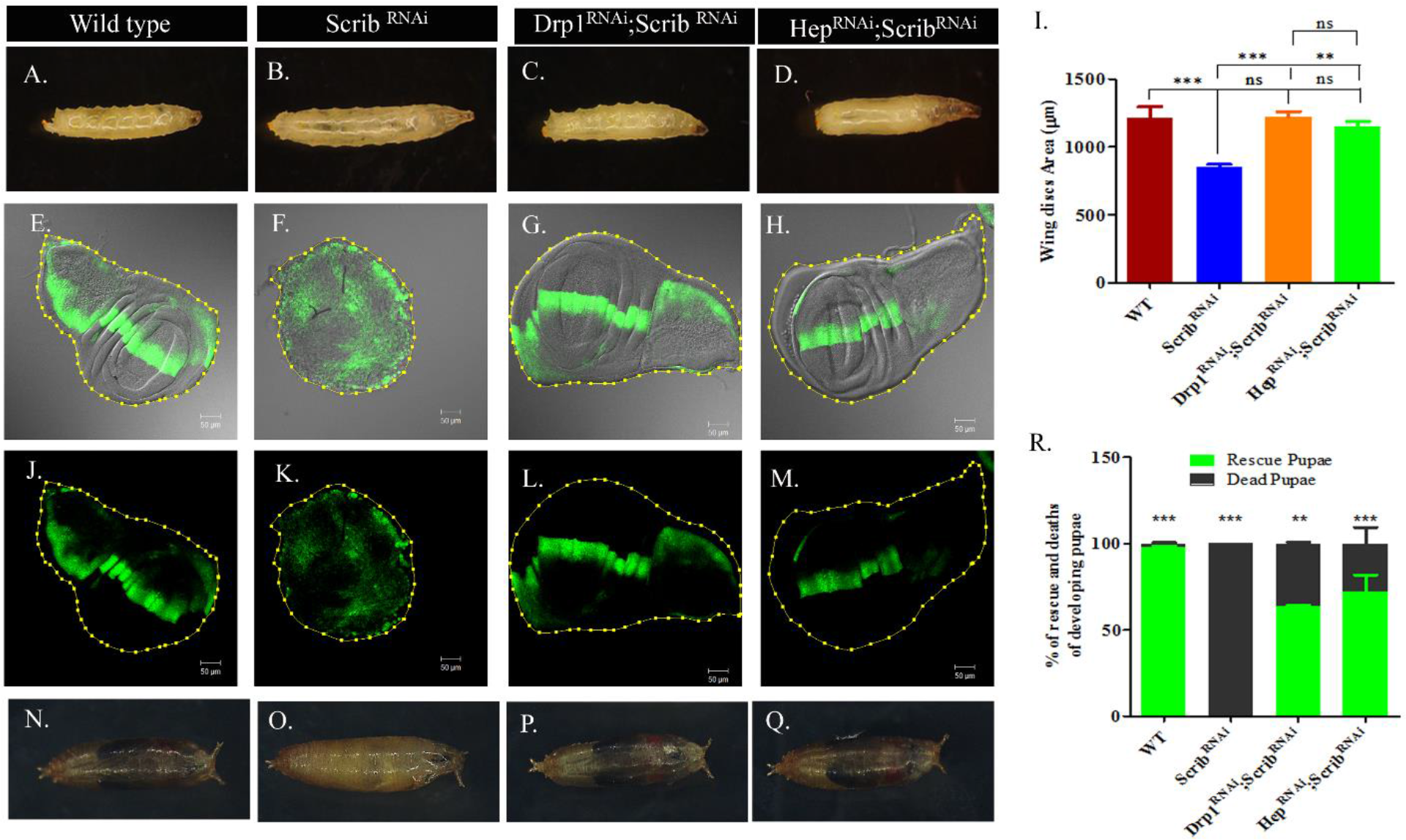
Rescue of *Scrib* loss induced phenotype by knockdown of *Drp1* and *JNK*. Mature 3^rd^ instar larvae from Wild type (A), *Scrib^RNAi^* (B), *Drp1^RNAi^; Scrib^RNAi^*(C) and *Hep^RNAi^;Scrib^RNAi^* (D). Note the enlarged larval size of *Scrib^RNAi^* (B) in comparison to wild type (A) and rescued phenotype (C-D). E-H: DIC + GFP images of normal WT wing discs (E) compared to *Scrib^RNAi^* (F) show neoplastic overgrowth whereas recused wing discs in *Drp1^RNAi^;Scrib^RNAi^*(G) *and Hep^RNAi^;Scrib^RNAi^* (H) show organized monolayer imaginal discs. Comparison of wing disc area in indicated genotype is shown (I, n=8). GFP-labeled Ptc-Gal4 expression (green) specifically in anterior to posterior region (J) of wing discs (UAS-GFP;Ptc-Gal4) whereas in *Scrib^RNAi^*wing discs (UAS-GFP;Ptc-Gal4>UAS-*Scrib^RNAi^*) show spreading of GFP labelled cells (K) and rescued wing discs UAS-GFP;Ptc-Gal4>*UAS-Drp1^RNAi^;UAS-Scrib^RNAi^*and UAS-GFP;Ptc- Gal4>UAS-*Hep^RNAi^;UAS-Scrib^RNAi^*show restoration of GFP expression pattern (L and M) as wild type. Wild type pupae show normal development inside the pupal case (N) while *Scrib^RNAi^* leads to 100% pupal death (O). The rescued pupae upon downregulation of *Drp1* (*Drp1^RNAi^; Scrib^RNAi^*) and *JNK (Hep^RNAi^; Scrib^RNAi^)* in *Scrib^RNAi^* background show complete fly development inside the pupal case (P and Q) as wild type. Histograms (R) represents percent of dead and live/ rescued pupae for indicated genotypes.

Loss of *Scrib* cell polarity regulator strongly promotes metastasis behavior of cancer cells [27]. To examine the contribution of *Drp1* mitochondrial fission protein and involvement of *JNK* signaling to prevent metastatic behavior in *Scrib* abrogated cells, we tagged Ptc-Gal4 driver line with GFP. The genetic crossing schemes are provided in supplementary data (Supplementary Information). In the control wing disc, the GFP expression pattern in Ptc-Gal4 is specifically in anterior posterior boundary region (Fig.1. J), whereas in *Scrib* knockdown there is a migration of cancer cells throughout the disc (Fig.1. K). In contrast, genetic knockdown of *Drp1* and Hep in *Scrib^RNAi^* background block cancer cell migration and restore the GFP expression pattern comparable to control wing disc (Fig.1.L and M). Taken together, these result indicate that inhibition of *Drp1* and *hep* in background of *Scrib^RNAi^* prevents cancer cell migration, clearly indicating the pioneer role of *Drp1* and *JNK* in prevention of metastasis.

Moreover, we found that loss of *Drp1/Hep* function suppresses pupal death upon *Scrib* loss and adult flies develop inside the pupal case. *Scrib* knockdown pupae exhibit 100% lethality at mid pupal stage (Fig.1 O) unlike wild type pupae that show 98% survival (Fig.1 N). Drp1 knockdown in *Scrib^RNAi^*background rescued 63% pupal lethality (Fig.1 P). However, *Hep* knockdown rescued 72% pupal lethality (Fig.1 Q). Histogram reveals percentage of rescue verses pupal death of developing pupae compared to wildtype (Fig.1 R).

### Knockdown of *Drp1 and JNK* signaling in *Scrib^RNAi^* background in wing imaginal discs restores to normal fly development and lifespan

The rescued flies of *UAS-Drp1^RNAi^; UAS-Scrib^RNAi^*and *UAS-Hep^RNAi^; UAS-Scrib^RNAi^* genotypes (Fig.2B and C, respectively) showed phenotypically normal morphology, wing development and reproduction comparable to wild type (A). Albeit, a slight difference was found in life span and wing venation pattern. The wildtype flies exhibited a mean lifespan of 84 days, the *Drp1^RNAi^*and *Hep^RNAi^* rescued flies showed a mean lifespan of 76 days, and 72 days, respectively.

**Figure 2.**
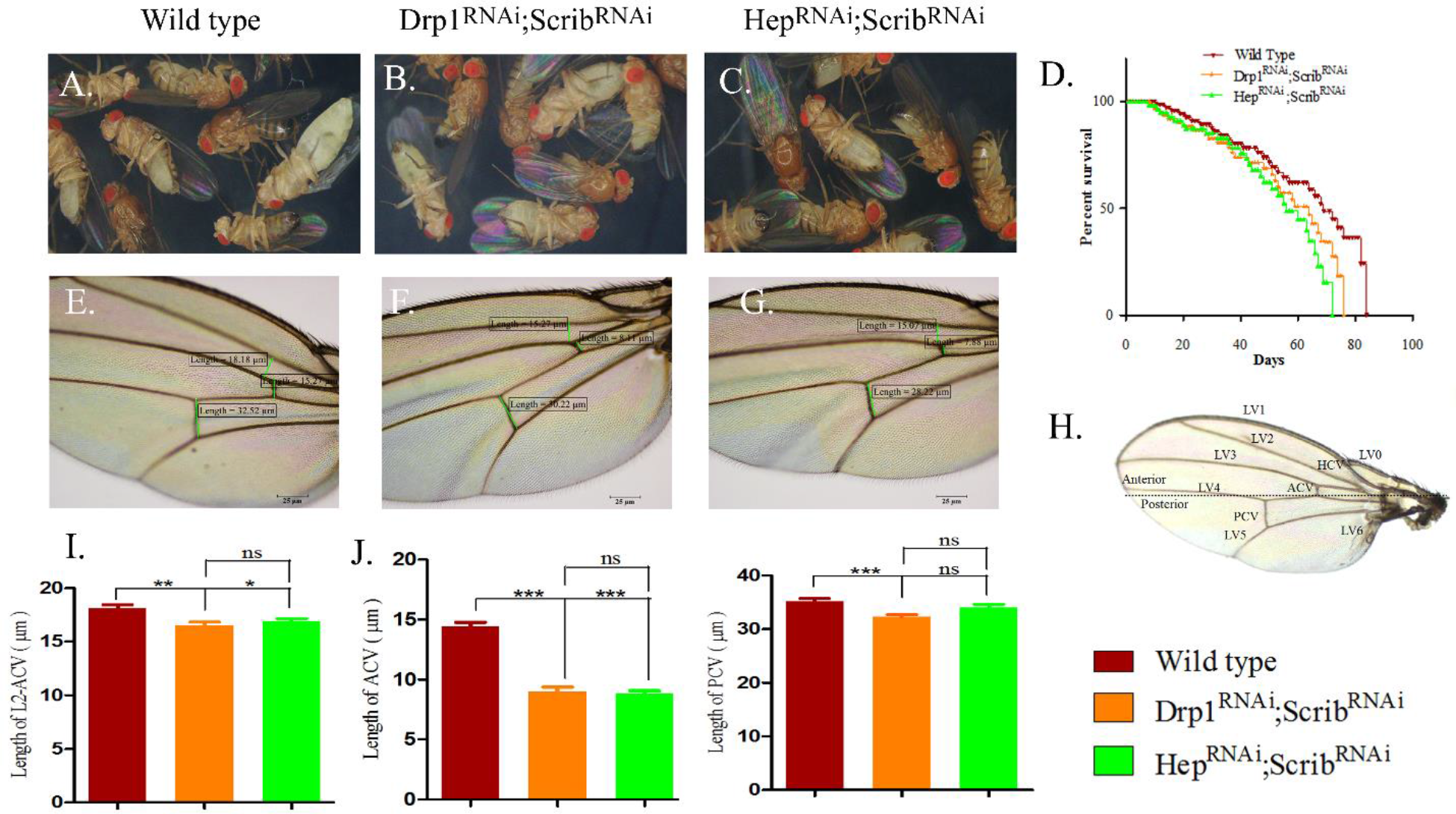
**Representing healthy escaper flies and adult wings venation pattern.**Panel A-C show mature healthy flies of Wild type, *Drp1^RNAi^; Scrib^RNAi^*and *Hep^RNAi^; Scrib^RNAi^.* Survival assay (D) showed life span of rescued flies (n=50). Examination of wing phenotype in wild type (E) showed normal wing morphology with regular arrangement of ACV, PCV and bristles while, *Drp1RNAi; ScribRNAi* (F) and *HepRNAi; ScribRNAi* (G) showed deformed wing vein pattern. Schematic diagram represents wildtype wing showing the location of longitudinal veins (L0-L6) and anterior cross vein (ACV) and posterior cross vein (PCV) (H). Quantification of wing morphology through measuring the length of L2-ACV (I), ACV (J) and PCV (K) between wild type and escaper flies (n=30) showed statistically significant difference in were analyzed by using one-way ANOVA (analysis of variances) followed by Bonferroni multiple comparison test (*p<0.05 **p<0.01 ***p<0.0001 ns p>0.005).

*Drosophila* adult wing venation is composed of two different types of veins, longitudinal veins (LV0-LV6) and the cross veins (ACV and PCV-anterior and posterior cross veins) [28]. According to latest research, *Scrib* protein regulates PCV in *Drosophila* wing to maintain epithelial morphogenesis [29,30]. Our findings suggested that upon genetic manipulation in *Scrib^RNAi^*, the length of different veins of rescued flies shortened (Fig. 2E-H). We report here significant changes in the length of L2-ACV (Fig. 2I), length of ACV that bridge L3-L4 (Fig. 2J) and PCV that bridge L4-L5 (Fig. 2K) upon loss of function of *Scrib* in combination of knockdown of *Drp1* and *Hep* gene in the wings of rescue flies compared to wildtype.

### *Drp1* inhibitor, *Mdivi-1* induces dose dependent inhibition of cancer cell migration

A small molecule quinazolinone derivative *Drp1* inhibitor*, mdivi-1* blocks the mitochondrial division by inhibiting the GTPase activity of the mitochondrial fission regulator, Drp1 [31]. We firstly show the effect of *mdivi-1* (**M**itochondrial **div**ision **i**nhibitor-1) on the maintenance of apico-basal polarity of cells upon loss of function of *Scrib*. In order to assess whether the inhibition of *Drp1* maintains the cell polarity, we screened a range of low to high concentrations (15µM to 500µM) of *mdivi-1* treatment on *Scrib* knockdown cancer bearing larvae (GFP; Ptc- GAL4>UAS-*Scrib^RNAi^*). Knockdown of *Scrib* in the wing discs induces disruption of cell shape and polarity, monolayer arrangement and patterning of wing disc leading to three dimensional overgrowth resulting in a deformed, tumorous disc showing a decrease in the wing disc area (Fig. 3B). Low doses (15µM and 30µM) of mdivi-1 fed-orally to tumor-bearing larvae do not show any phenotypic change on the tumor size (data not shown), whereas 60µM and 120µM mdivi treatment showed an increase in the area of the disc (Fig. 3C and D). However, at 250µM and 500µM concentrations of mdivi, the monolayer epithelial wing imaginal disc morphology is restored (Fig. 3E and F).

**Fig 3.**
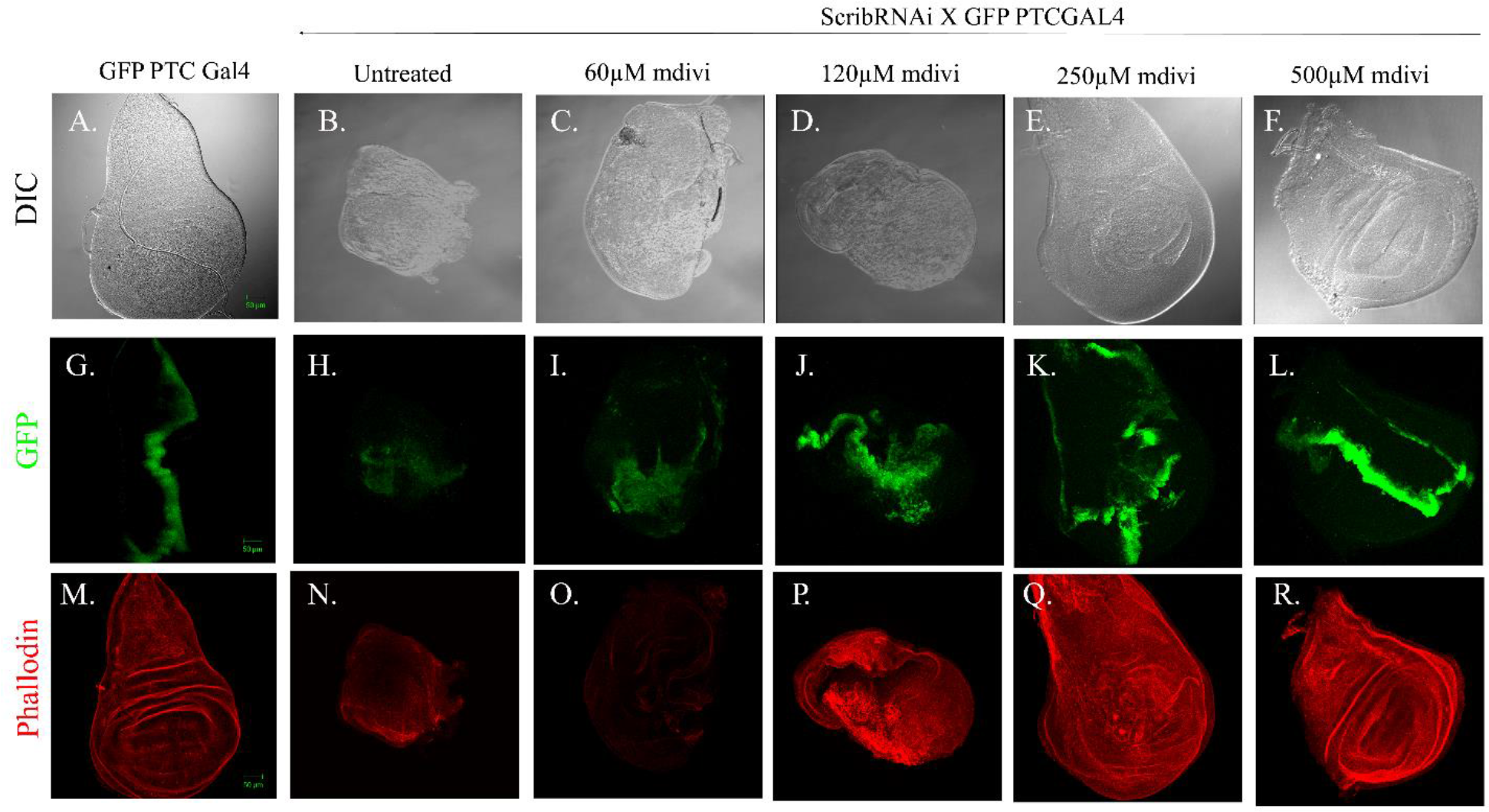
Effect of different doses of *Mdivi-1*treatment on *Scrib^RNAi^* cell migration. DIC images showing the untreated control wing imaginal discs (A) in comparison to the deformed, tumorous disc (B), and gradual restoration of normal morphology upon increase in the dosage of *Mdivi-1* treatment (C-F). The control wing disc shows GFP expression pattern specifically in the anterior to posterior boundary (G) whereas *Scrib^RNAi^*shows the spreading of GFP positive cells (H). *Mdivi-1* treated *Scrib^RNAi^*larvae (I-L) show gradual restoration of GFP expression pattern. Rhodamine Phalloidin staining of wing discs representing proper F-actin arrangement in WT (M) as compared to disruption of actin in *Scrib^RNAi^* wing disc (N), while restoration of actin cytoskeleton and inhibition of neoplastic overgrowth can be seen in mdivi-1 treated wing imaginal discs (O-R).

Further, the *mdivi-1,*was also effective against cancer cell migration and minimized the metastasis in *Scrib* knockdown wing imaginal disc. In third instar wildtype larvae (GFP; Ptc- GAL4), GFP expression pattern is confined specifically to a single strip in the mid region of the monolayer epithelial wing disc (Fig. 3G). This pattern is lost in the Ptc-Gal4 driven *Scrib^RNAi^*(GFP; Ptc-GAL4>UAS-*Scrib^RNAi^*), and the GFP-tagged cancer cell migration is found beyond the Ptc expressing domain in the tumorous disc (Fig. 3H). Oral administration of low doses (15µM and 30µM) of mdivi-1,does not show alteration in the GFP expression pattern compared to the untreated tumorous wing disc (data not shown). At 60µM, there is an increase in the number of GFP expressing cells, but no change is found in the expression pattern of GFP (Fig. 3I). 120µM mdivi concentration shows effective response and partially restores the expression pattern of GFP (Fig. 3J). 250µM *mdivi-1*furtherinhibited the migration of GFP tagged *Scrib^RNAi^* cells (Fig. 3K). 500µM of mdivi-1 blocked the migration of GFP-tagged *Scrib^RNAi^* cells into the surrounding area and completely restored the GFP expression pattern comparable to wildtype, establishing it to be the effective concentration required for the rescue (Fig. 3L).

Moreover, we also observed the effect of mdivi-1 on the cytoskeleton arrangement of F-actin counterstained with rhodamine phallodin (Fig. 3M-S). The wildtype disc shows a proper arrangement of actin (Fig. 3M), while loss of function of *Scrib* alters the distribution of actin filaments in the tumorous wing disc (Fig. 3N). Low doses (15µM and 30µM) of mdivi-1 treatment showed no changes in actin arrangement. However, 60µM and 120µM mdivi-1 treatment partially rescued the arrangement of actin (Fig. 3O-P). Further, 250µM *mdivi-1* treatment shows significant improvement in the cytoskeleton arrangement of F-actin (Fig. 3Q).

*Mdivi-1* high dose (500µM) restored the proper arrangement of phallodin labeled actin, confirming it to be the effective dose for the rescue of *Scrib* knockdown tumorous wing disc (Fig. 3R).

Collectively, the above data revealed that a high dose (500µM) of mdivi-1 treatment decreases cell proliferation, inhibits cell migration, restores F-actin cytoskeleton arrangement and maintains apico-basal cell polarity. We show here that the dose-dependent effect of mdivi-1 to inhibits cell proliferation and cell migration in the tumorous wing disc.

### *Scrib* regulates polarity of cells through the Drp1-JNK signaling pathway

The downregulation of *Scrib* activity contributed to elevated *Drp1-JNK* activation, which leads to exacerbation of the apico-basal cell polarity defects in *Drosophila*. We and others demonstrated that loss of *Scrib*function leads to loss of cell polarity with absolute pupal death during early stage of fly pupal development [14]. Our novel findings suggested that, loss of *Drp1* and *JNK* upstream regulator, *Hep* gene function in *Scrib* knockdown background results in restoration of cells polarity and recovery from early death of *Drosophila* pupae. To explore the molecular mechanistic action of *Drp1-JNK* signaling in regulation of cell polarity and metastasis upon loss of *Scrib*, we measured the mRNA expression level using qRT-PCR and protein expression level using immunostaining and western blotting analysis. Immunostaining revealed that *Scrib* knockdown tumorous disc shows upregulation of Drp1, PJNK and MMP1 expression as compared to wildtype and rescued wing discs (Fig. 4A-L).

**Fig. 4.**
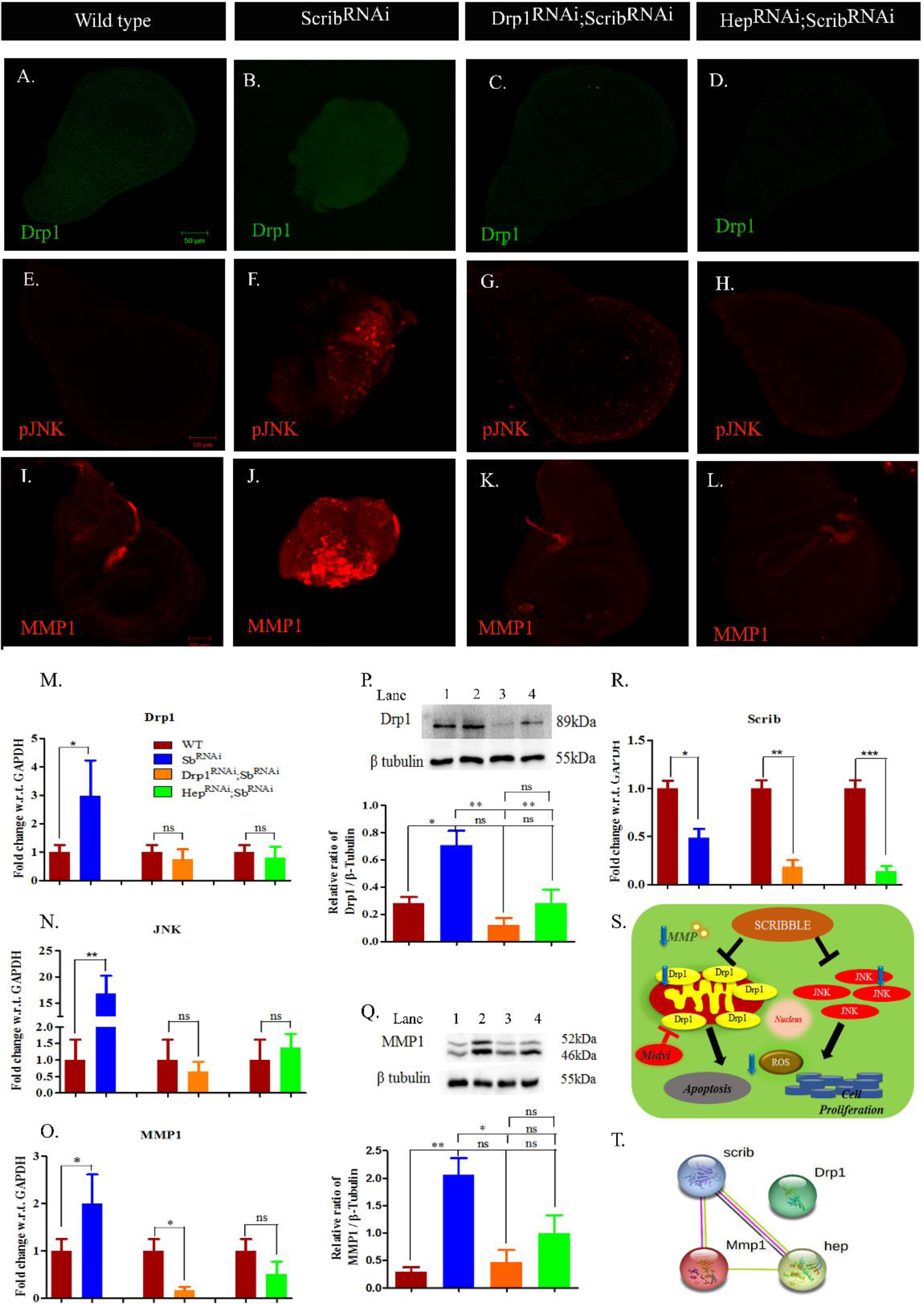
**Panel shows immunostaining, Real-time PCR and Western blot analysis for Scrib, Drp1, pJNK and MMP1 in *Drosophila*.**A-L Wing disc from Wild type, *Scrib^RNAi^*, and *Drp1RNAi; Scrib^RNAi^* and *Hep^RNAi^; Scrib^RNAi^* rescue genotypes showing Drp1 (A-D), pJNK (E-H) and MMP1 (I-L) staining. Wing disc from *Scrib^RNAi^* showed significant upregulation of Drp1 (green, B), phospho-JNK (red, F) and MMP1 (red, J) expression as compared to wild type and rescued wing discs (*Drp1^RNAi^; Scrib^RNAi^* and *Hep^RNAi^; Scrib^RNAi^*). Real-time PCR (M-O, R) and western blot (P and Q) for Drp1, JNK and MMP1 also revealed the upregulation of Drp1, JNK and MMP1 expression in *Scrib* knockdown as compared to wild type (A) and rescue genotypes. Lane 1, 2, 3 and 4 represents the protein band intensity for Wild type, Scrib^RNAi^, *Drp1RNAi; Scrib^RNAi^* and *Hep^RNAi^; Scrib^RNAi^* respectively (S) A schematic model showing *Scrib* inhibits the gene expression of *Drp1* and *JNK* to retained the polarity of the cells and also to regulate cell proliferation, apoptosis and maintenance of ROS levels. (T) PPI network of Drp1 with Scrib, JNK and MMP1. Statistically significant differences were analyzed between wildtype and tested group of indicated genotype by using one-way ANOVA (analysis of variances) followed by Bonferroni multiple comparison test and two-tailed student’s t test for real-time PCR (*p<0.05 **p<0.01 ***p<0.0001 ns p>0.005).

Further, to validate the immunostaining results, we performed qRT-PCR to measure gene expression level of *Drp1, JNK* and *MMP1* in *Scrib* knockdown compared to wildtype and rescue pupae. The mRNA transcript level of *Drp1*is 2.97 fold upregulate in *Scrib* knockdown while 0.73 and 0.81 fold downregulated in *Drp1^RNAi^; Scrib^RNAi^* and *Hep^RNAi^; Scrib^RNAi^,* respectively (Fig. 4M). Further, the mRNA transcript level of *JNK* is 16.87 fold upregulated in *Scrib* knockdown while downregulation in rescue pupae shows 0.64 fold in *Drp1^RNAi^; Scrib^RNAi^*and 1.37 fold in *Hep^RNAi^; Scrib^RNAi^* (Fig. 4N). Whilst, the mRNA transcript level of *MMP1* is 1.99 fold upregulated in *Scrib* knockdown while 0.16 and 0.50 fold downregulated in *Drp1^RNAi^; Scrib^RNAi^* and *Hep^RNAi^; Scrib^RNAi^* in rescue genotype (Fig. 4O). No significant difference was found in *Drp1^RNAi^; Scrib^RNAi^* and *Hep^RNAi^; Scrib^RNAi^* compared to wildtype. A similar result was found in western blot assays where protein expression of Drp1 (Fig. 4P) and MMP1 (Fig. 4Q) was upregulated in *Scrib* knockdown while in the rescue pupae, *Drp1^RNAi^; Scrib^RNAi^* and *Hep^RNAi^; Scrib^RNAi^* the protein level is downregulated.

Further, to confirm the knockdown efficiency of Ptc-gal4>*Scrib^RNAi^*upon co-induction of *Drp1^RNAi^*and*Hep^RNAi^*, being driven by the same Gal4, we performed RT-PCR for the *Scrib* gene. The RT-PCR gene expression analysis revealed that Ptc-Gal4>UAS-*Scrib^RNAi^*line could efficiently silence the *Scrib* gene despite the co-expression of *Drp1^RNAi^* and *Hep^RNAi^*transgenes.

The mRNA transcript level of *Scrib* is 0.48 fold downregulated in *Scrib* knockdown and 0.17 fold and 0.13 fold downregulated the rescue genotypes, *Drp1^RNAi^; Scrib^RNAi^* and *Hep^RNAi^; Scrib^RNAi^*, respectively compared to wild type (Fig. 4R).

The schematic model represents the summary of molecular studies. It shows that *Scrib* positively regulates the expression of *Drp1* and, *Hep* genes to maintain the polarity of cell and suppresses the cancer cell migration by depletion of MMP1 gene expression (Fig. 4S). Using STRING database, a protein-protein interaction (PPI) network diagram was automatically generated for Scrib, Drp1, Hep and MMP1 proteins.The protein Scrib, has an experimental link with JNK upstream regulator, Hep and metastasis marker, MMP1 protein whereas, Drp1 protein, did not show any interaction. The Scrib protein is co-expressed with Hep protein. So, this in-silico study clearly reveals that *Drp1* has no previous experimental connection with polarity regulator Scrib protein, metastasis regulator MMP1 protein and JNK upstream regulator, Hep protein. Our results for the first time show the experimental connection between Drp1 with Scrib protein and also with JNK and MMP1protein, to maintain the polarity of the cells (Fig. 4T).

### Knockdown of *Drp1* and *Hep* in *Scrib^RNAi^* background reduces ROS production and oxidative stress markers

ROS plays a vital role in regulation of neoplastic tumor growth upon loss of *Scrib*[32]. Therefore, we examined the effect of *Drp1* depletion and *JNK* inactivation on ROS production upon loss of *Scrib*. To detect in-vivo intracellular ROS production in *Scrib knockdown* wing imaginal disc, we used H2DCFDA (2’,7’-dichlorodihydro fluorescein diacetate) staining, a cell membrane permeable indicator used for detection of ROS. In-vivo cellular ROS production in wildtype shows weak signals (Fig.5A) whereas in *Scrib KD* wing imaginal discs show very strong green fluorescent signal suggested excess ROS production in tumor bearing wing imaginal discs (Fig.5B). Furthermore, rescued wing imaginal discs *Drp1^RNAi^; Scrib^RNAi^*and *Hep^RNAi^; Scrib^RNAi^* show inhibition of ROS production as revealed by faint green fluorescent signal as compared to wildtype (Fig.5C and 5D).

**Fig 5.**
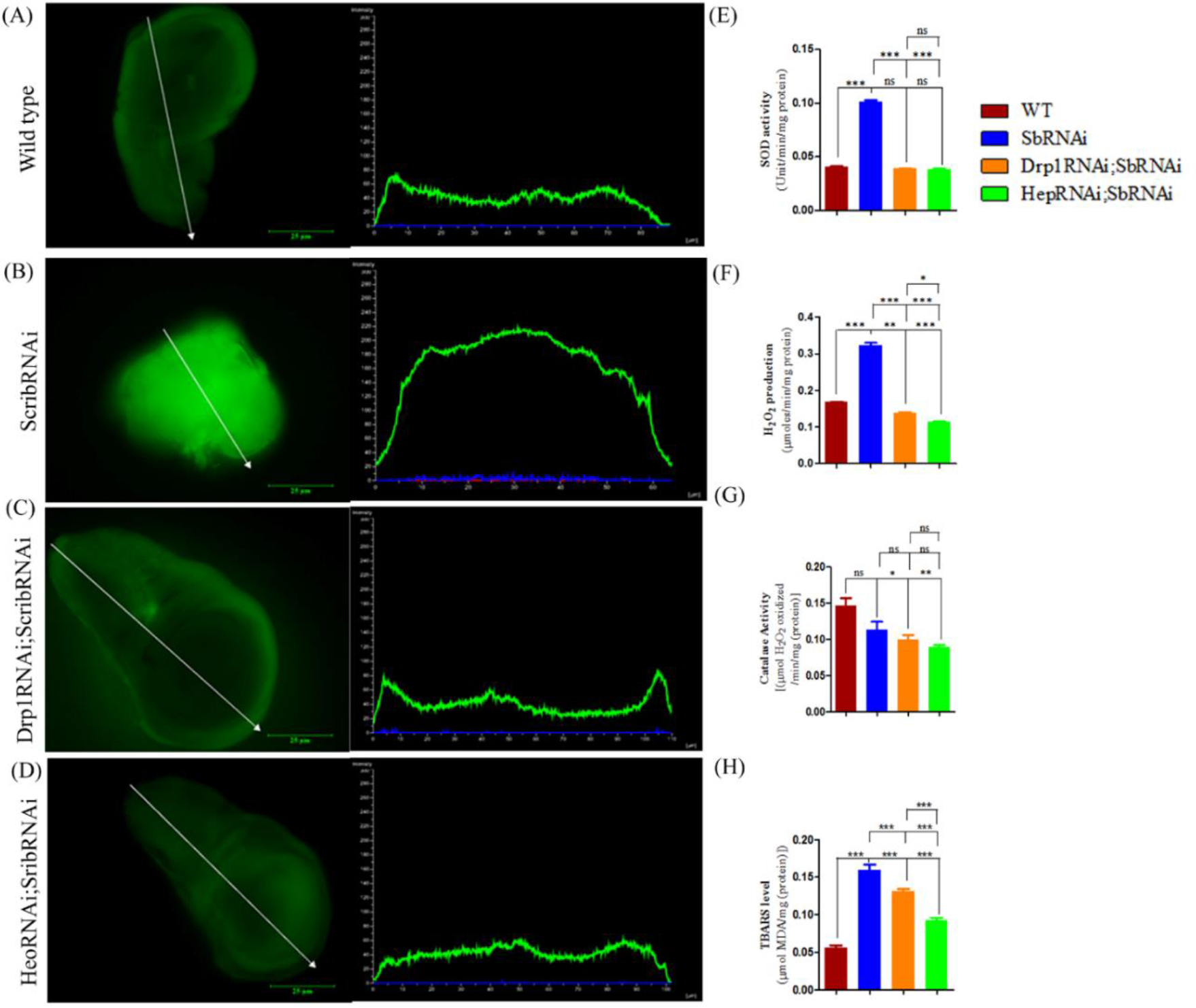
H2DCFDA staining and biochemical assays to detect *in situ* ROS and oxidative stress. *In situ* ROS detection using 2,7dichlorodihydrofluorescein diacetate (H2DCFDA) staining (green), and imaging by the fluorescence microscope (A-D). The wild type (A) wing discs show low level in situ cellular ROS as compared to tumorous wing imaginal discs (B) while in rescued wing discs (*Drp1^RNAi^; Scrib^RNAi^* and *Hep^RNAi^; Scrib^RNAi^*) ROS production is comparable to control (C and D). The Intensity line graphs showed total ROS production in wild type, *Scrib^RNAi^* and rescued wing discs along the white arrow drawn by NIS-Elements BR 4.3 software. SOD activity (E), H2O2 production (F), Catalase activity (G) and TBARS level (H) measured in *Scrib^RNAi^*and rescued group compare to wild type. The statistical significance was analyzed using one-way ANOVA (analysis of variances) followed by Bonferroni multiple comparison test (*p<0.05 **p<0.01 ***p<0.0001 ns p>0.005).

Further, biochemical assays such as SOD (superoxide dismutase) activity assay, H2O2 (hydrogen peroxide) production assay, Catalase activity assay and LPO (lipid peroxidation) assay, were also performed for detection of ROS production. In tumor cells, elevated mitochondrial ROS production is altered by aberrant expression of antioxidant enzymes, like superoxide dismutase, catalase and excess production of lipid peroxidation (LPO) and hydrogen peroxide (H2O2), a characteristic marker for oxidative stress [33]. We determined the activities of antioxidant enzymes in wildtype compared to *Scrib* knockdown and genetic rescue pupae to check the status of oxidative stress (OS). The *Scrib* KD showed 2.49-fold higher SOD activity while the rescued genotypes *Drp1^RNAi^; Scrib^RNAi^* and *Hep^RNAi^; Scrib^RNAi^* showed reduced SOD activity,1.04 and 0.95 fold respectively, almost comparable (Fig.5E).

Superoxide dismutase, catalyzes the superoxide radical to hydrogen peroxide, H2O2. So, we further evaluated the status of H2O2 production. *Scrib^RNAi^*, displayed significant upregulation of H2O2 compared to wildtype and rescue pupae (Fig.5F). The excess H2O2 production is further catalyzed by antioxidant catalase enzymes, which converts H2O2 to water and molecular oxygen [34]. *Scrib^RNAi^* pupae displayed reduced catalase activity compare to wildtype and rescued pupae (Fig.5G). Moreover, to determine the effect of excessive ROS production in the context of lipid peroxidation, TBARS assays was performed. The pink color end product was measured using multimode plate reader at absorbance of 532nm and 600nm [35]. The lipid peroxidation marker, TBRAS level shows significant elevation in *Scrib* KD cells whereas, in rescued pupae it showed reduction of LPO in form of reduce in TBRAS levels as compare to wildtype (Fig.5H).

### Knockdown of *Drp1 and JNK* signaling in loss of *Scrib^RNAi^*regulates the balance between cell proliferation and apoptosis to maintain the mitochondrial morphology

To investigate the role of Drp1 in maintenance of apico-basal cell polarity by regulating the Drp1- mediated mitochondrial fission in *Scrib* knockdown cells, we examined mitochondrial distribution in control, Scrib knockdown, Drp-1 knockdown and JNK knockdown under Scrib knockdown backgrounds in wing imaginal discs. We used an outer mitochondrial membrane tagged mitocherrythat showed healthy and defective mitochondria in wildtype and *Scrib^RNAi^*respectively (Fig. 6A and B). *Scrib* knockdown wing disc contained severely defective, aggregated, fragmented and randomly distributed mitochondrial throughout the tumorous wing disc, unlike the wild type where an orderly mitochondrial arrangement is seen across anterior to posterior region of wing discs. Strikingly, when *Drp1* and *Hep* were knockdown in *Scrib^RNAi^*cells, *UAS-Drp1^RNAi^; UAS-Scrib^RNAi^* and *UAS-Hep^RNAi^; UAS-Scrib^RNAi^*, the mitochondrial structure and arrangement was restored as wildtype, specifically in anterior posterior boundary region (Fig.6.C and D; Cross scheme provided in supplementary data). The magnified images are shown in Fig.6 A’-D’.

**Fig 6.**
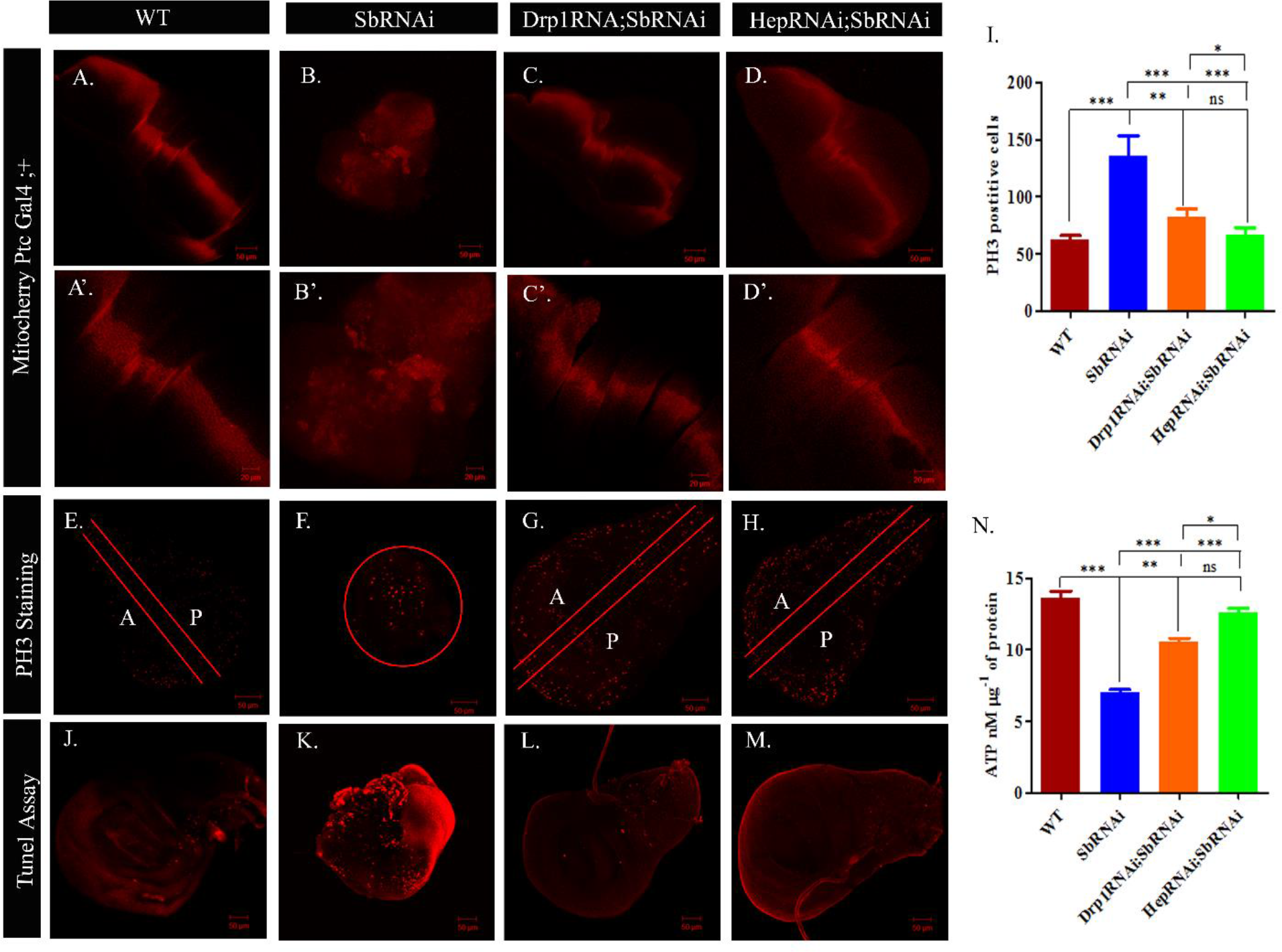
Confocal images representing mitochondrial distribution (A-D’), anti-Ser10 phosphorylated histone (PH3) staining (E-H) and Tunel assay (J-M) in wing imaginal discs. (A, A’) Wild type discs (genotype mitocherry Ptc-Gal4;+) showed well-arranged mitochondria, specifically in anterior and posterior region of wing imaginal discs whereas *Scrib^RNAi^*discs showed distorted and aggregated arrangement of mitochondria throughout the tumorous wing discs (B, B’). The rescued wing discs (C, C’ and D, D’) showed similar arrangement as Wildtype. The magnified confocal projection showed rescued flies wing imaginal disc shows maintenance of similar mitochondrial morphology as wild type compared with *Scrib^RNAi^* tumorous discs (Fig.5.A’ and D’). Anti-PH3 staining in wild type (E) wing imaginal discs showed less PH3 positive cells across anterior to posterior boundary whereas in *Scrib^RNAi^* cells (F) shows overlap and aggregate PH3 stained + cells while in rescued wing discs shows less PH3 positive cells in A/P boundary region of wing imaginal discs (G and H). A, indicate anterior region and P, indicate posterior region of *Drosophila* wing imaginal disc. The histogram shows the significant increasein PH3 positive cells in tumor disc as compare to wildtype and rescued wing disc (I). Panel (J-M) shows tunel assay of wild type (J) and *Scrib^RNAi^* wing imaginal discs (K). Number of tunel positive cell (red) are greater in *Scrib^RNAi^* cells compared to wild type while the rescued wing imaginal discs (Drp-1^RNAi^ and Hep^RNAi^) show less tunel positive cells (L and M). Histogram showing the reduction of ATP levels in *Scrib^RNAi^* while significant restoration of ATP levelswere observed in rescued pupae as compared to *Scrib^RNAi^* pupae (N).

Earlier well documented studies show that loss of *Scrib* promotes excessive cell proliferation and evades apoptosis [26,36]. Here, our results show that upon loss of Drp1 and hep in background of *Scrib^RNAi^*, control the cell proliferation and regulates the apoptosis as revealed by PH3 staining and tunel assays respectively. The abnormal and uncontrolled growth of *Scrib* knockdown cells induces excessive cell proliferation. Therefore, we firstly assess the cell proliferation status across the anterior to posterior boundary upon knockdown of *Drp1* in *Scrib^RNAi^*background by assessing expression of Phospho-histone PH3 staining, marker for proliferating cells (indicated by red color) (Fig. 6.E-H). In the wildtype wing imaginal discs, very few PH3 positive cells were observed in anterior-posterior (A/P) boundary region of the discs (Fig.6.E). However, in tumorous disc larger number of PH3 positive cells enrichedaround apparent A/P region which is encircled in red and spread all over the wing disc (Fig.6.F) while lesser number of PH3 positive cells present in A/P boundary region of rescued wing imaginal disc upon depletion of Drp1 and JNK inactivation in background of *Scrib* knockdown (Fig.6.G and H). Further, quantification of PH3 positive cells also shows significant larger number of proliferating cells in *Scrib^RNAi^*as compared to wildtype while rescued wing imaginal discs showed less number of PH3 positive cells (Fig.6.I). These results suggested that depletion of Drp1 positively regulates the over proliferation of the cancer cells upon knockdown of *Scrib*. Similarly, upon knockdown of JNKupstream regulator, Hep gene also positively control the cell proliferation in background of *Scrib^RNAi^* for maintaining the apico-basal polarity of the cells.

In cancer, there is a competition between cell proliferation and apoptosis which plays major role in survival of cancer cells [37]. So, to determine whether loss of cell polarity upon knockdown of *Scrib* induces apoptosis, we examined third instar larval wing imaginal discs which are subjected to the tunel assay for marking the apoptotic cells. There is a striking increase in the number of apoptotic cells in tumorous wing disc as compared to wildtype (Fig.6.J and K). Furthermore, the knockdown of *Drp1* and *Hep* gene in background of *Scrib^RNAi^*shows decrease in number of tunel positive cells in rescued wing imaginal discs (Fig.6.L and M), suggesting that Drp1-mediated mitochondrial fission contributes in apoptosis regulation upon loss of *Scrib* gene to maintain the cell polarity.

Mitochondria are double membrane bound dynamic cell organelles that are power house for energy production and also prime source for ROS production [38]. In several diseases, the overproduction of ROS is associated with a decrease in ATP synthesis due to mitochondrial dysfunction which fail to maintain cellular energy [39]. Therefore, it is important to measure the ATP levels upon *Drp1* depletion in background of loss of *Scrib*. Notably, our results clearly show that unlike wild type, overproduction of ROS leads to significantly reduced the ATP production in *Scrib* knockdown suggesting mitochondria dysfunction (Fig.6.N). Furthermore, the depletion of *Drp1* and *Hep* significantly restores the ATP production (Fig.6.N).

## Discussion

*Drosophila* third instar larval wing imaginal discs are monolayered epithelial cells that develop and differentiate into adult appendages, the wings, and most frequently used for modeling cancer [8], including our current study. *Scrib*, the apico-basal polarity regulator gene of epithelial cells is necessary for maintenance of epithelial homeostasis [40]. According to earlier research findings, the Scrib, a lateral membrane associated protein found at cell-cell junction (adherens junction-AJs). Scrib is mislocalized to the cytoplasm of cancer cells upon loss of function mutation in Scrib complex proteins [41–43]. The *loss of function* mutation of *Scrib* leads to development of neoplastic tumor [27,44]. Earlier reports from our laboratory showed larval wing tissue specific knockdown of *Scrib* developed neoplastic tumors with increased larval size and die as giant pupae during early stages of pupal development without differentiation [14]. The tumorous wing imaginal discs have three dimensional multilayered proliferation of cells rather than highly organized monolayered flattened epithelial cells as in wild type discs. The molecular mechanisms of polarity regulation upon loss of *Scrib* remains elusive. In recent years, the role of mitochondrial-dynamics regulators has been well investigated in cancer progression and metastasis [14, 45]. Cancer cells show differential expression of mitochondrial dynamics regulator genes leading to enhanced fission-fusion events, causing disruption of mitochondrial morphology. The fusion-fission events are critical for maintenance of mitochondrial dynamics and polarity aspects of cells. The mitochondrial fission and fusion events are critically affected in several diseases including cancer and metastasis conditions. The Drp1 is emerging as key regulator of fission events and JNK is also emerging as key component in Drp1 mediated regulation of mitochondrial dynamics. Although the role of mitochondria in several types of cancer and neurodegenerative diseases have been well investigated, but the role of Drp1-JNK mediated mitochondrial fission event in regulation of cell polarity maintenance remains unexplored.

To further explore the molecular mechanistic action of *Scrib* on mitochondria dynamics, the double transgenic fly lines (*UAS-Drp1^RNAi^; UAS-Scrib^RNAi^*and *UAS-Hep^RNAi^; UAS-Scrib^RNAi^*) were created to investigate the role of Drp1 and JNK in regulation of cancer progression and metastasis. To test our hypothesis that the upregulation *Drp1* in *Scrib* knockdown tumorous tissues is responsible for neoplastic tumor development, we genetically downregulated *Drp1* expression levels using *UAS-Drp1^RNAi^*, in the background of *UAS-Scrib^RNAi^*<Ptc-Gal4 (referred as a *UAS-Drp1^RNAi^; UAS-Scrib^RNAi^*), which resulted in significant reduction in tumor growth and complete rescue leading to development of healthy *Drosophila* flies. The same phenotype was also observed by downregulation of JNK upstream regulator, Hep, in the background of *UAS- Scrib^RNAi^*<Ptc-Gal4 (referred as a *UAS-Hep^RNAi^; UAS-Scrib^RNAi^*). We genetically modulated *Drp1/Hep* expression in *Scrib* knockdown background using *RNAi* approach and show the rescue of cancer phenotypes, resulting in healthy adult escaper flies (Fig. 1). We further show for the first time that the *Drp1* depletion in *Scrib* knockdown cells rescue from extended larval development and larval overgrowth phenotypes.

To further confirm the effect of *Drp1^RNAi^* and *Hep^RNAi^* in cancer bearing larvae, we allowed third instar larvae to mature into adult flies and examined the morphological alteration, if any, and reproductive aspects compared to wild type counterparts. The escaper flies (*UAS-Drp1^RNAi^; UAS- Scrib^RNAi^*and*UAS-Hep^RNAi^; UAS-Scrib^RNAi^*) were healthy except exhibiting a minor differences in wing venation pattern of wing but the reproduction and lifespan were similar to wild type (Fig. 2). The phenotypic, biochemical and molecular changes upon knockdown of *Drp1* expression encourages us to exploit the chemical approach and examine the effect of mdivi-1, a *Drp1* inhibitor on the *Drp1* mediated mitochondrial division. For this, we assessed the dose dependent effect of mdivi-1 on cancer bearing *scrib* knockdown larvae. The mechanistic action of mdivi-1 on the GTPase activity of *Drp1* suggest that it inhibits the mitochondrial fission and positively regulates the polarity of *scrib*knockdown cells. We observed the restoration of wing imaginal disc morphology by mdivi-1 to maximum extent but it could not rescue from pupal lethality phenotype. The fact that mdivi-1 restores wing disc morphology of Scrib^RNAi^ suggests Drp-1role in cancer progression and metastasis. Mdivi-1 could not rescue from metastasis at pupal development. Inhibition of *Drp1* mediated restoration of mitochondrial fission is considered as an efficacious therapeutic targets for the prevention of cancer. Further studies are required to evaluate how exactly mdivi-1 and Drp-1 play role in regulation of mitochondrial division and regulating the cell polarity in the absence of Scibble (Fig. 3).

Our molecular studies suggest that *Scrib* regulates polarity of cells through the *JNK-Drp1* mediated mitochondrial fission event. The downregulation of *Scrib* contributed to *Drp1* activation, which led to migration of *Drp1* from the cytoplasm onto the surface of the mitochondria, indicative of mitochondrial fission activation. Earlier reports from our laboratory clearly showed that the *Scrib^RNAi^* knockdown leads to upregulation of *Drp1*which was rescued through *HepRNAi*[14]. In order to check whether Drp-1 downregulation under *Scrib^RNAi^*knockdown background can rescue the loss of cell polarity and pupal lethal phenotype, we further genetically modified the expression of *Drp1* in *Scrib^RNAi^* background using *Drp1^RNAi^* lines, which restored cell polarity as evident by the restoration of wing discs morphology and eventually rescued the pupal death phenotype. Wevalidated the downregulation of *Drp1* by Immunostaining, qRT-PCR, western blotting analysis. Although it is clear that *Drp1* regulate JNK pathway [46,47] but its role in *Scrib* mediated neoplastic tumor development is completely unknown. We went onto show that *Drp1* is also key target of *JNK* downstream regulator, *Hep* gene that contributes to neoplastic tumor growth. To test this hypothesis, we genetically knocked-down the *Hep* gene using *Hep^RNAi^* line in *Scrib^RNAi^*background. The loss of *Drp1*- mediated fission event downregulates the *JNK* expression while loss of *Hep* gene downregulates the *Drp1* expression in *Scrib* knockdown cells. Immunostaining, RT-PCR and western blotting data revealed that genetic manipulation using RNAi lines leads to silencing of the *Drp1/Hep*gene expression in *Scrib^RNAi^* background. Overall, our results for the first time provide evidence that *Drp1* and *JNK* activation leads to tumor progression and *Drp1-JNK* inhibition by genetic and chemical inhibition means reverse cancer progression and metastasis as a downstream mechanism of *Scrib* knockdown cells gives new insight into the mechanistic action of *Scrib* in cancer cells. In summary, it is concluded from these studies that reduced *Drp1*-*JNK* mediated fission events in tumor cells help maintain the apico-basal polarity of the cells. *Drp1* knockdown inhibits mitochondrial fission event and maintain mitochondrial homeostasis in *Scrib* knockdown cells. Moreover, down regulation of *Drp1/Hep* genes under *Scrib^RNAi^* background can also suppress malignant behavior of metastatic secondary tumors which is a major causative event of cancer related deaths [48]. The elevated levels of *MMP1* expression during metastasis window period and its rescue by down regulation of MMP1 were evaluated by using qRT-PCR for RNA expression, immunostaining for *in situ* MMP1 localization and western bolt analysis for protein quantification clearly demonstrated that the downregulation of Drp1 and JNK play role in altering MMP1 to restore normalcy. Our data clear shows, *Scrib* regulates mitochondrial fission via the *JNK-Drp1* mediated pathways to control cancer progression and metastasis. This novel involvement of *Scrib-JNK-Drp1* pathway in regulating the cell proliferation, maintaining the polarity of cells through modulating the dynamics of mitochondria in cancer cells can be considered as strategic therapeutic target for several advance stage cancers (Fig. 4).

Several types of human cancer have been reported to be associated with mitochondrial dysfunction and ROS production [47]. Mitochondrial dysfunction is a prime source of overproduction of ROS resulting in oxidative stress [49]. The imbalance between ROS production and impaired enzymatic activity of antioxidant agents leads to mitochondrial dysfunction in cancer. *Scrib* knockdown elevated activity of SOD, H2O2 and LPO, reduced catalase activity while *Drp1* and *JNK* downregulation under *Scrib* knockdown background showed reduced activity of ROS. The reduced oxidative stress by *Drp1/JNK* downregulation in *Scrib* knockdown background indicate the role of *Drp1* and *JNK* in activating the oxidative stress while Scribble counteract Drp-1 and JNK to maintains the oxidative parameters and polarity of the cells in an optimal manner (Fig. 5). It is intriguing to note that downregulation of Drp-1 and JNK restoring the cell polarity and rescuing lethality phenotype despite loss of Scribble could be possibly due to blocking mitochondrial fission events, which require further studies.

Overall, *Scrib* plays a role in maintaining the cell polarity and cell proliferation regulation by blocking the *Drp1* and *JNK* which regulate mitochondrial dynamics, via maintaining the mitochondrial membrane polarity, morphology and oxidative stress. Mitochondria, a dynamic double membrane-bound cell organelles, play crucial role in ATP production, recently emerged as hub for Drp-1 mediated mitochondrial fission protein interactome connective pathways for various diseases [50]. Our results clearly show that upon knockdown of *Scrib* leads to enhanced Drp-1 mediated disruption of mitochondrial dynamics, which induces enhanced ROS production and cause reduced ATP production leading to increased cell proliferation and metastasis. On other hand, Drp1 and JNK downregulation together with *Scrib^RNAi^* suppress ROS production, enhances ATP production, maintains cell polarity and restores normal fly development. These findings suggest that mitochondrial fission regulator, Drp1is key modulator of altered mitochondrial dynamics upon loss of *Scrib* through altering the ROS production, ATP generation, excess cell proliferation and metastasis. It is well documented that Drp-1 plays is a central role in triggering the mitochondrial fission mediated metastasis and apoptosis [51]. Our findings also show Drp1 and JNK depletion enhance cell death phenomena in *Scrib* abrogated cells thereby restoring normal development. More has to be probed into the mechanistic aspects of Drp-1-JNK-Scrib related cell regulation in normal development versus cancer progression and metastasis.

## Conclusions

The Scrib-JNK-Drp1 pathway play unique role in regulating planer cell polarity (PCP) of epithelial cells. Dysfunction of above pathway is observed in loss of *Scrib*. Our findings demonstrated for the first time that *Drp1*-knockdown and *mdivi* treatment markedly suppressed excessive cell proliferation and regulates the apico-basal cell polarity loss induced by knockdown of the *Scrib*. We also found that *Hep* inactivation attenuates the *Drp1* mediated mitochondrial dysfunction and regulate the PCP in the *Scrib* knockdown induced tumor cells. The alteration in mitochondrial fission regulator, Drp1 provide important clues towards prevention and therapeutic intervention for the cancer drug discovery.

## Acknowledgments

This work was supported by a research grant (P07/598) awarded to SS by the Science and Engineering Research Board, India and ICMR-Senior Research Fellowship (ICMR-SRF) to JS. Authors are highly grateful to IoE (Institution of Eminence) for financial support under incentive grant and SATHI for central facility. The authors would like to acknowledge ISLS-BHU and for Confocal Microscopy, Nanodrop, Real time PCR and other routine facilities.

## Supplementary Information

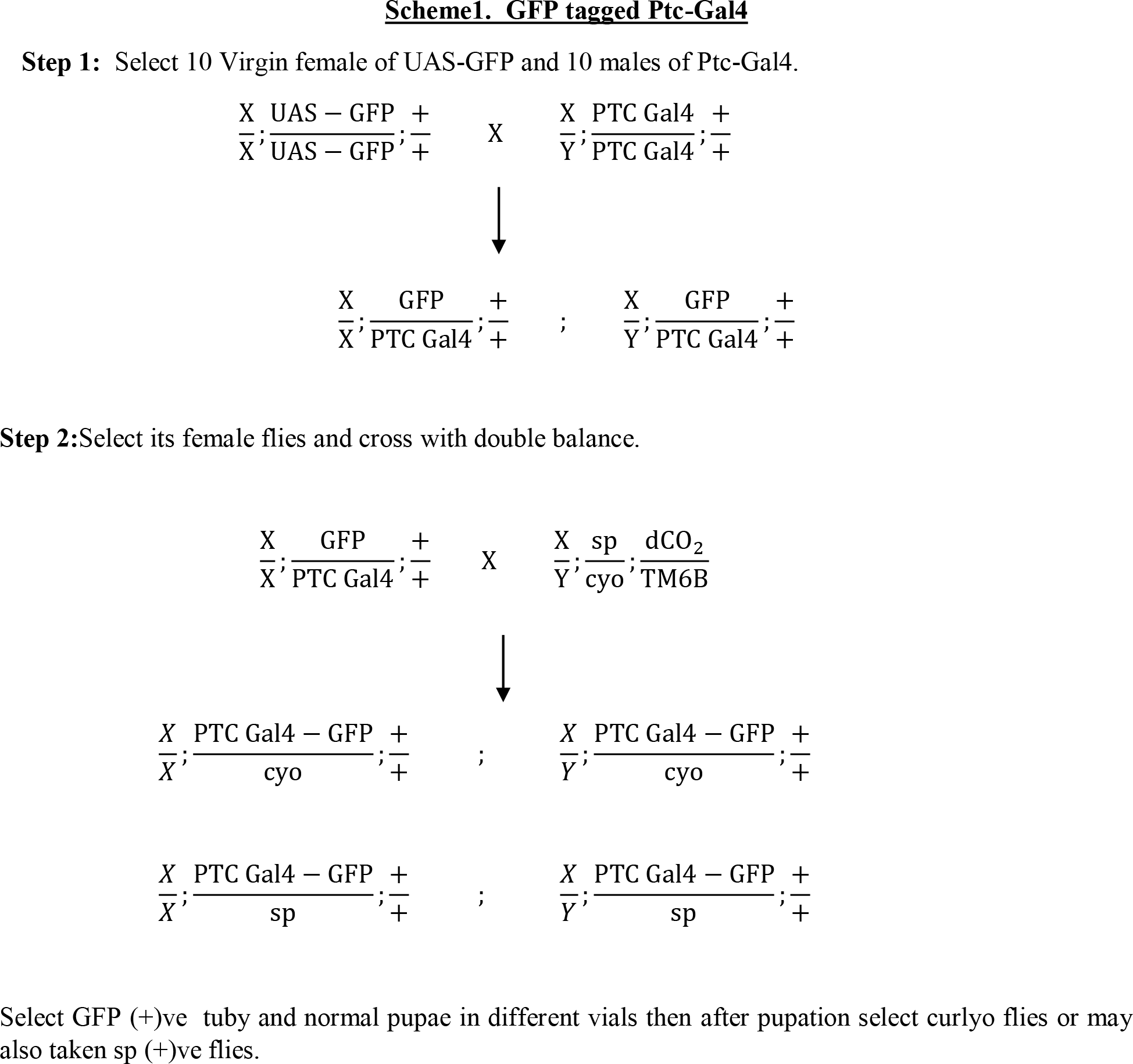
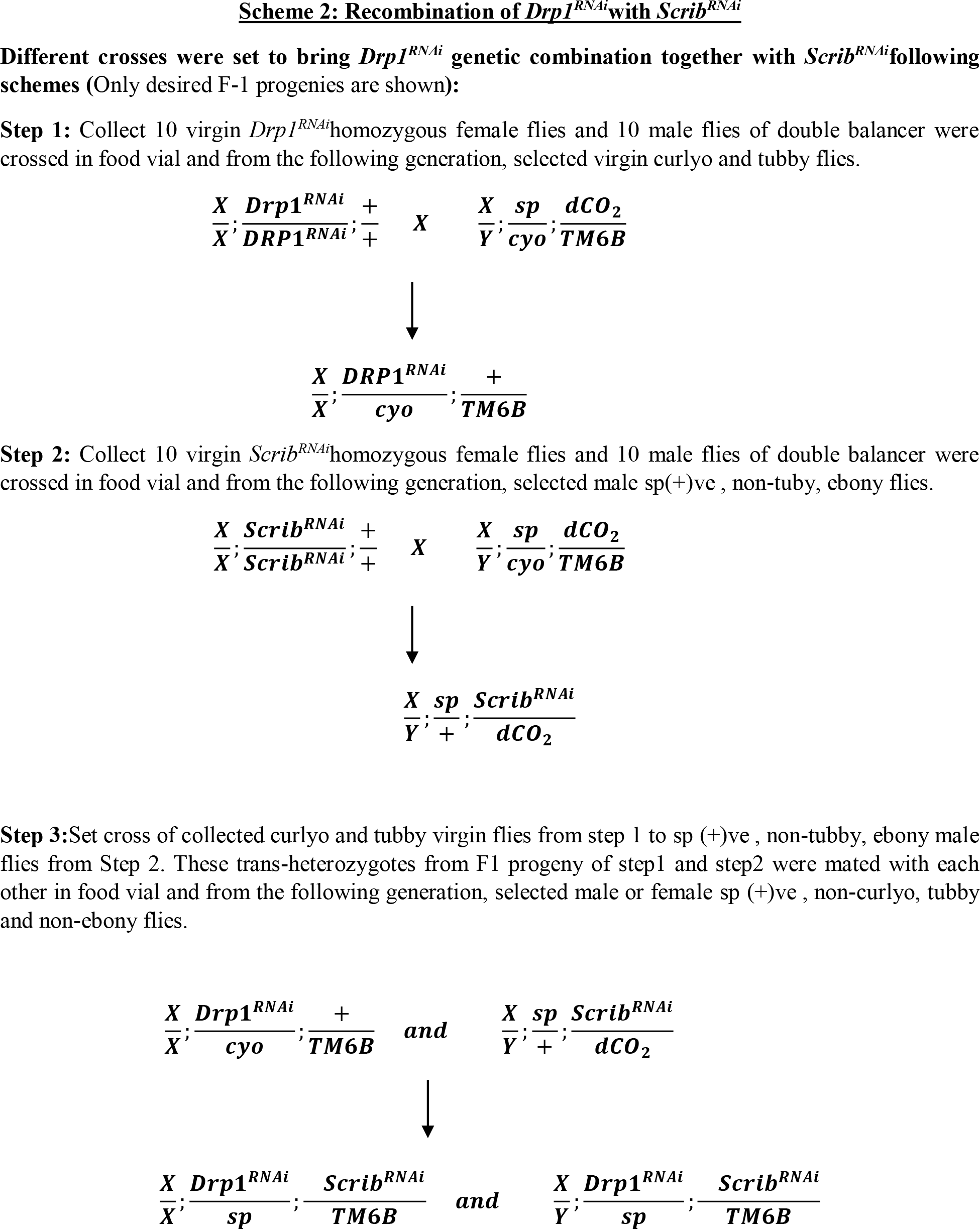
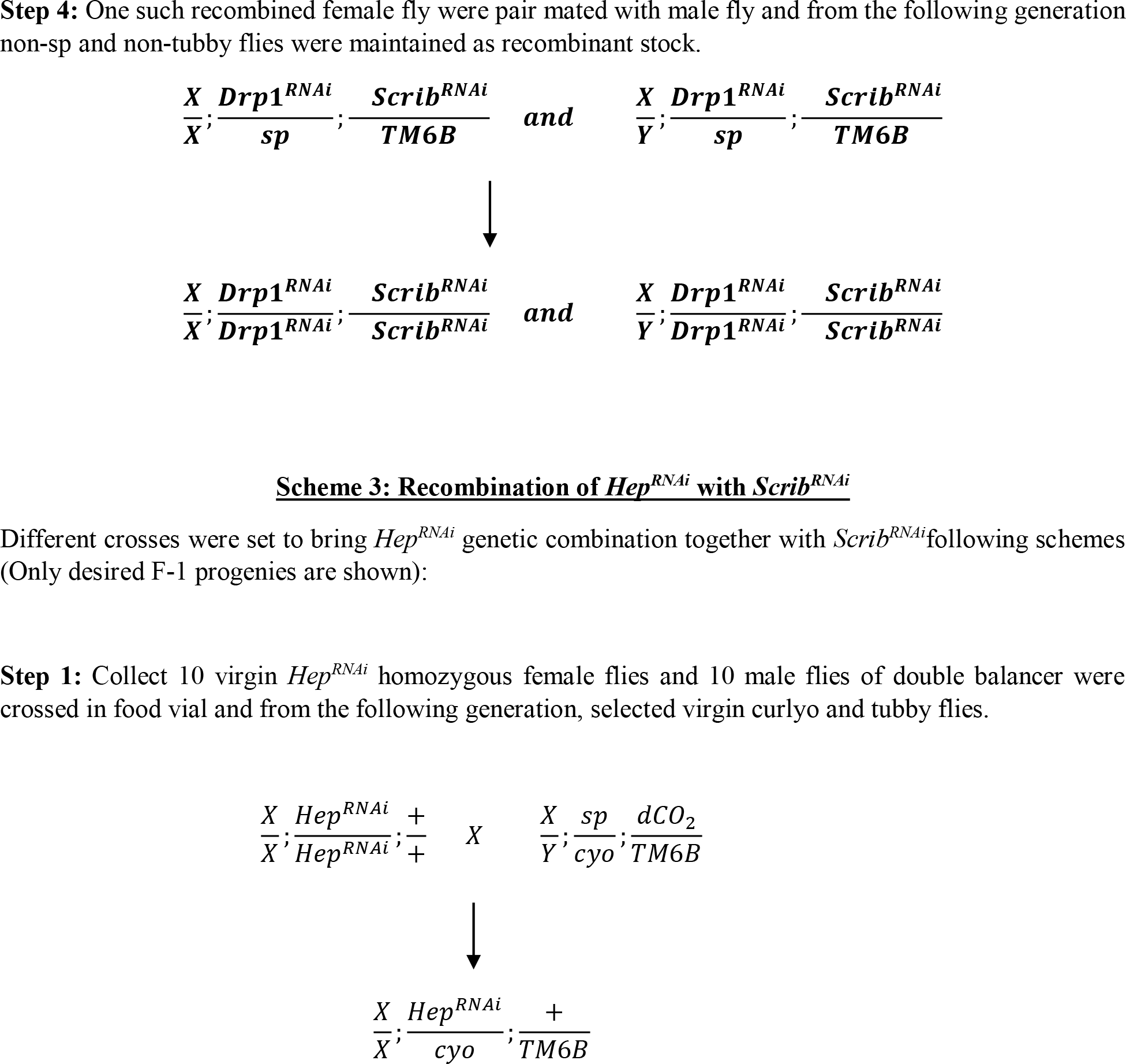
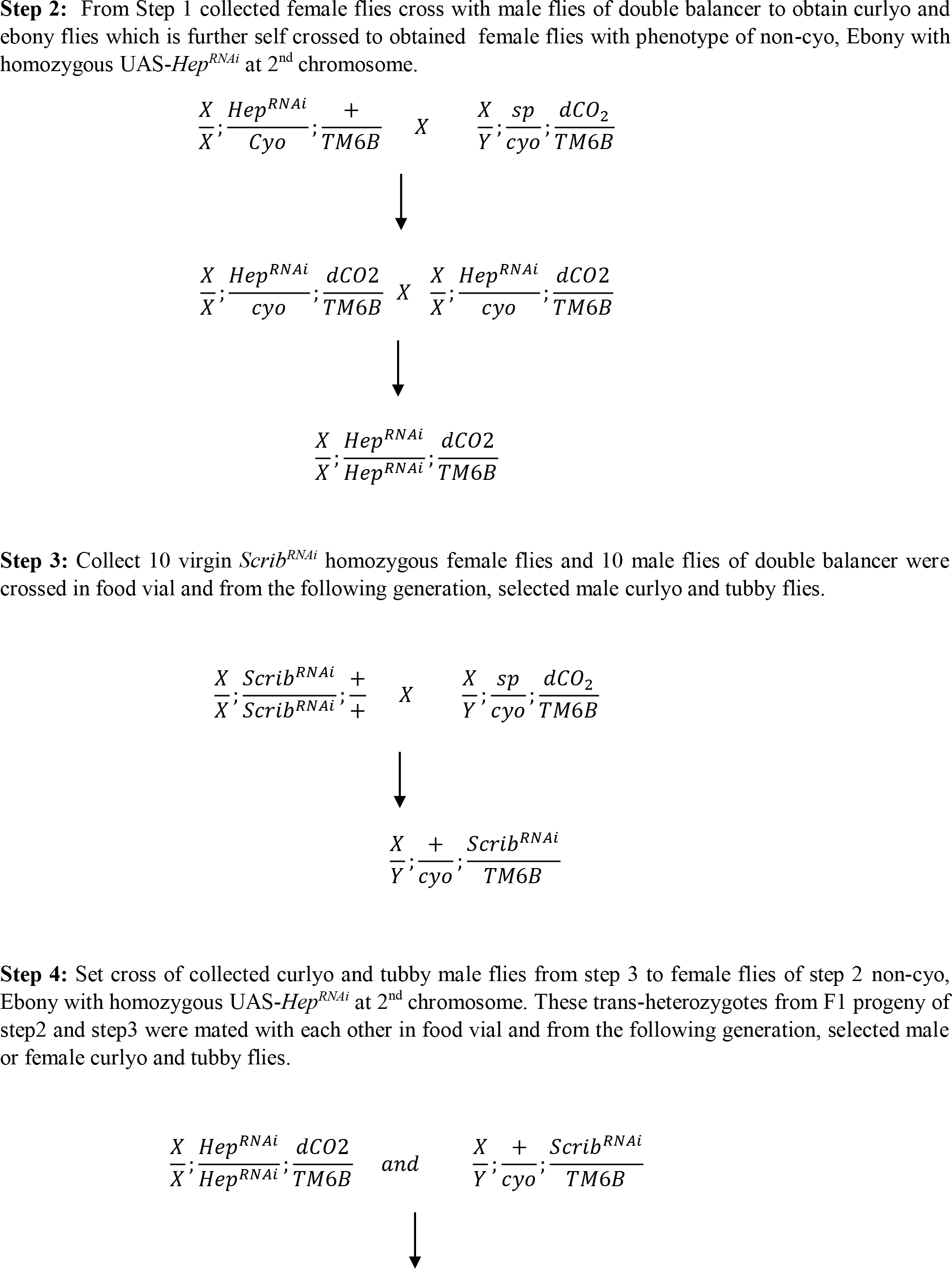
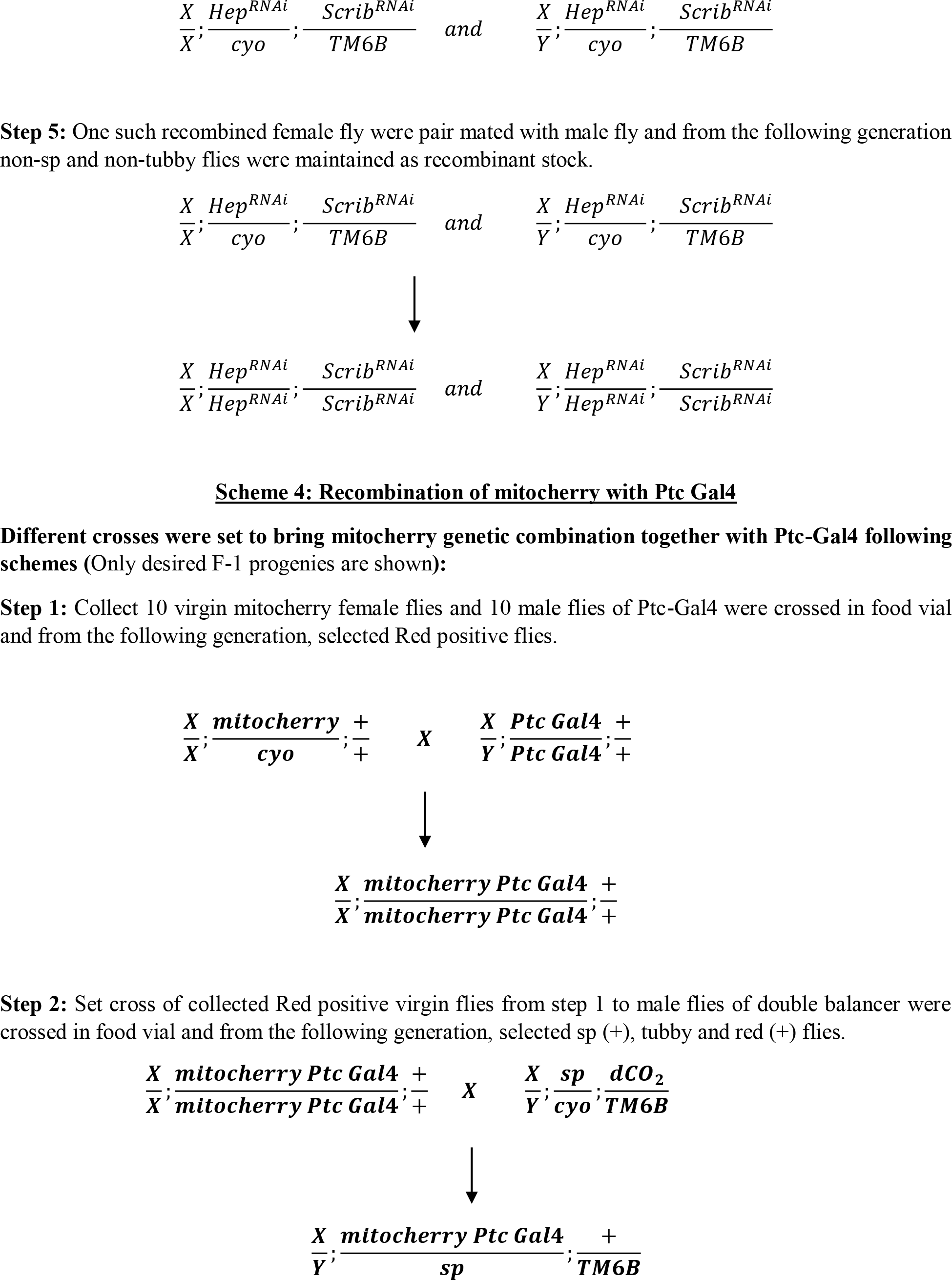
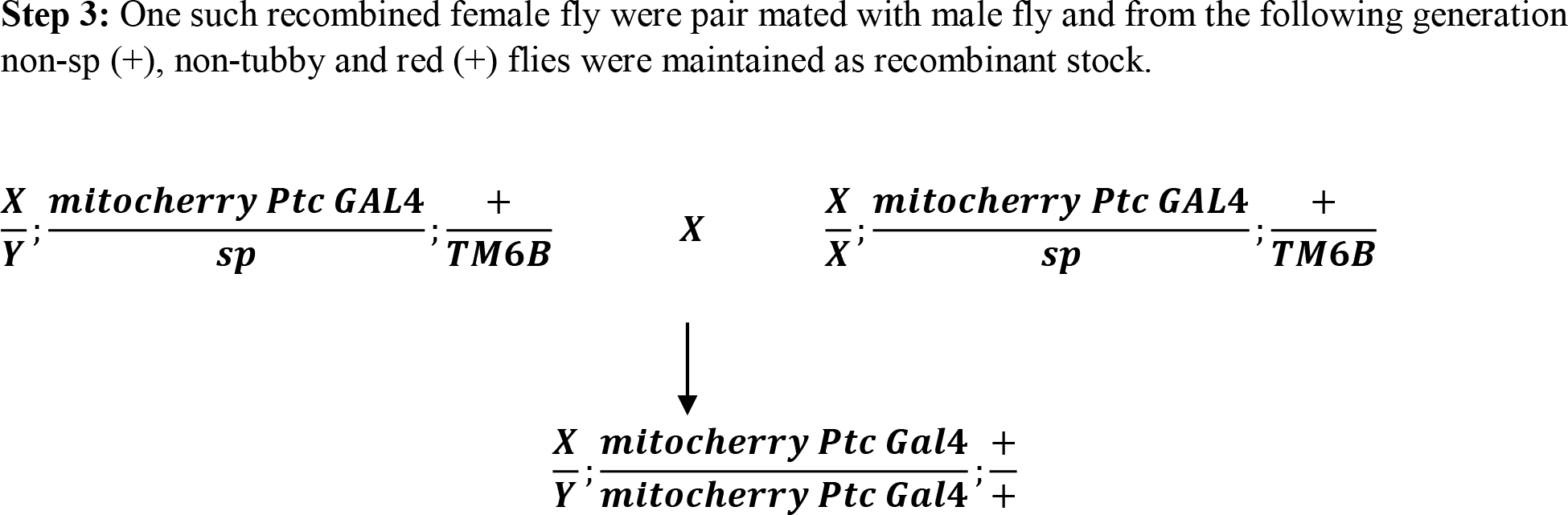

